# Nodulation Trio in *Medicago truncatula:* Unveiling the Overlapping Roles of MtLYK2, MtLYK3, and MtLYK2bis

**DOI:** 10.1101/2024.02.19.577609

**Authors:** Yaohua Li, Yanwen Zhao, Ziang Yan, Ru Dong, Haixiang Yu, Hui Zhu, Yangrong Cao

**Affiliations:** National Key Lab of Agricultural Microbiology, Hubei Hongshan Laboratory, Huazhong Agricultural University, Wuhan, 430070, China

**Keywords:** *Medicago truncatula*, Nod factor receptor, nodulation, nodules

## Abstract

The perception of rhizobial nodulation factors (NF) by plant NF receptors is crucial for mediating legume nodulation. In Medicago truncatula nodulation, a proposed two-receptor model designates MtLYK3 as an entry receptor, while other LYK proteins, with unidentified functions, may serve as signaling receptors. Here, we generated the single, double, and triple mutants for M. truncatula MtLYK2, MtLYK3, and MtLYK2bis using CRISPR/Cas technology and examined their roles in nodulation. Our findings suggest that all three LYKs possess redundant functions in Medicago nodulation, with their distinct contributions attributed to varying transcription patterns.Our findings also suggest that a one-receptor model with multiple specificities that mediates both signaling and entry responses is more suitable in M. truncatula.

---

The legume-rhizobial symbiosis is a distinctive model in plant-microbe interactions. Despite being foreign invaders, rhizobia are acknowledged as “friends” by compatible host plants, leading to mutual endosymbiosis [1]. Crucial to the establishment of this endosymbiosis is the recognition of rhizobial nodulation factor (NF) by a hetero-receptor complex involving two Lysin-Motif receptor Kinases (LYKs) in plants, namely NF receptor1 (LjNFR1) and LjNFR5 [2-4], as observed in *Lotus japonicus*. The knockout of either *LjNFR1* or *LjNFR5* results in a complete loss of rhizobial symbiosis [2, 4]. However, the situation is different in *Medicago truncatula*[5], where three *LYK* genes-*MtLYK3, MtLYK2*, and *MtLYK2bis* are identified as orthologs of *LjNFR1* (Fig. 1B). *M. truncatula spp. truncatula* cv Jemalong A17 (referred to as A17) contains *MtLYK2* and *MtLYK3*, while *M. truncatula spp. tricycla* R108 (referred to as R108) has all three LYKs. In A17, the *MtLYK3* knockout mutant, *hcl (hair curling)*, loses the ability to form a shepherd’s crook and infection thread but keeps some NF-signaling responses, including calcium spiking and expression of some symbiotic genes [6, 7]. These data support a two-receptor model in Medicago nodulation: MtLYK3 may function as an entry receptor, while the other LYK protein with uncharacterized functions may serve as signaling receptors [5, 7, 8]. These findings pose an intriguing question about how these three LYKs coordinate to regulate the rhizobial response. Here, we generated the single, double, and triple mutants for *M. truncatula MtLYK2, MtLYK3*, and *MtLYK2bis* using CRISPR/Cas technology and examined their roles in nodulation. Our findings suggest that individual LYK proteins can fully participate in the symbiotic process of NF recognition.

**Fig. 1.**
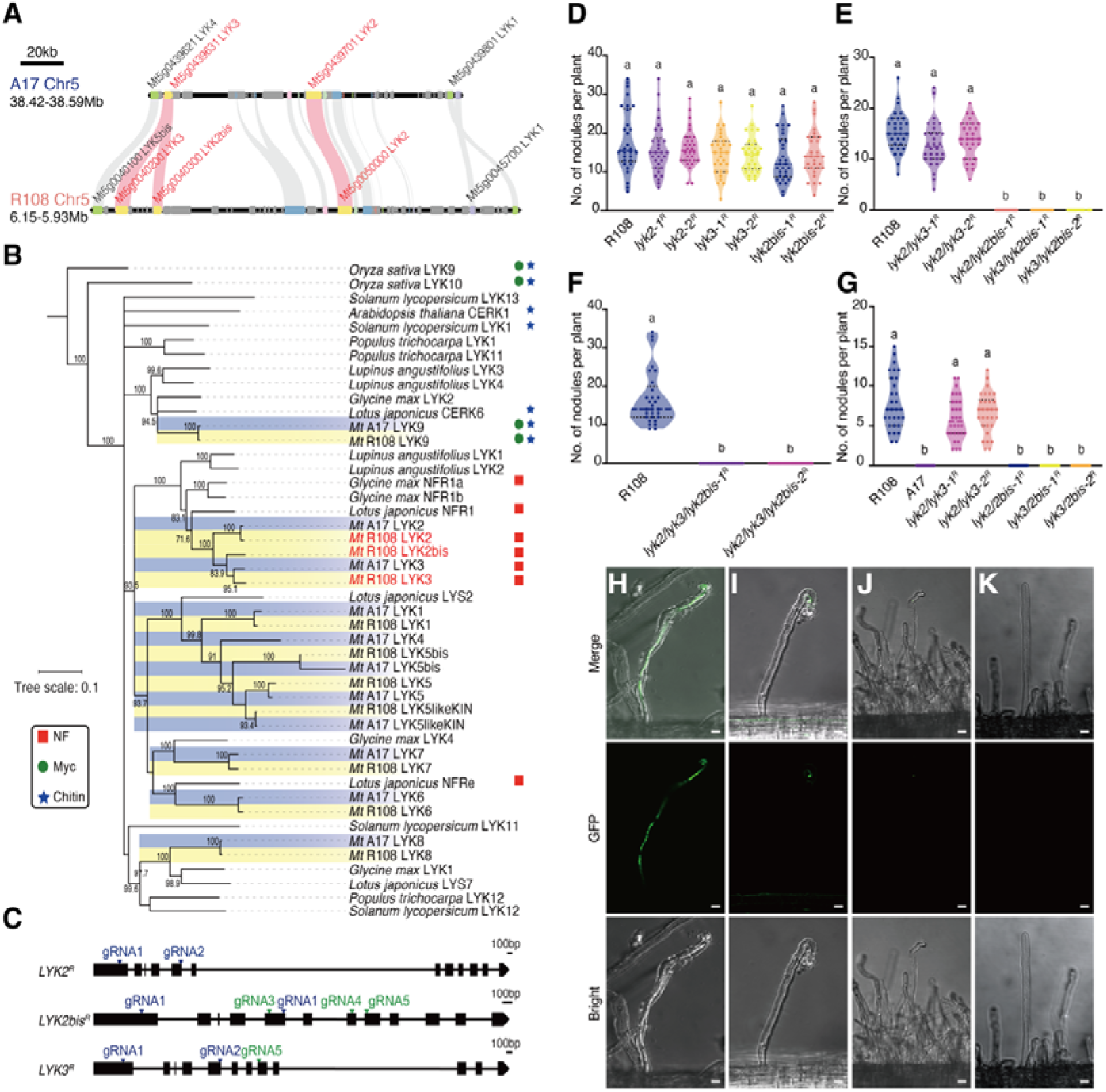
Phenotypic analysis of *Medicago truncatula* Mutants. (A) Microsynteny analysis reveals partial homology between the *Medicago truncatula* A17 and R108 genomes in the 5th chromosome region. The syntenic blocks encompass specific genes from the LYK-I orthogroup gene family. (B) Phylogenetic reconstruction of the LYK-I orthogroup, encompassing known LCO receptors, is conducted using 46 sequences derived from 8 species. Two subgroups were identified in *M. truncatula* were identified to *Mt*A17 (bule color) and *Mt*R108 (yellow color), respectively, and the focal proteins were marked in red font. Proteins with a known function in nodulation, mycorrhisation and/or chitin signalling were marked using red squares, green circles and blue stars for labelling, respectively.The numerical values on the branches of the evolutionary tree represent support values, indicating the confidence level for each branch. A list of species and protein sequence can be found in Dataset S1. (C) Schematic representation of *LYK2, LYK2bis* and *LYK3* of R108, showing the structure of the genomes and the gRNA target sites in their exon. Exon sequences are shown in black blocks; Intron sequences are represented by lines. (D,E and F) Analysis of nodulation in R108 mutants. Number of nodules at 21 days post-inoculation with S. meliloti 2011 for single mutants (D), double mutants (E), and triple mutants (F). (G)nodules number formed on different plants 21 d post inoculation with *S. meliloti* 2011 *nodf/nodl* in double mutants. Analysis was conducted on 30 plants for each genotype/inoculum. Lowercase letters indicate significant differences (ANOVA, Tukey, P < 0.05). (H, I, J and K) Development of epidermal infection threads in *M. truncatula* root hairs at 4 days post-inoculation. (H)Typical infection by *S. meliloti* 2011 (expressing GFP; green) of a root hair on WT R108 plants. (I)In the *lyk2bis/lyk3-1* mutant plants, hair curling and infection sites were observed. (J)In the *lyk2/lyk3-1* mutant plants, only infection sites were found. (K)In the *lyk2/lyk2bis/3-1* mutant plants, no hair deformation, hair curling, or infection sites were observed.Merged-field images are displayed on the upper panel, GFP-field images on the middle panel, and bright-field images on the lower panel. (Scale bar, 20 μm)

In R108, the three LYK genes, *MtLYK3, MtLYK2*, and *MtLYK2bis*, are located in the same cluster on chromosome 5 (Fig. 1A). This clustered arrangement complicates the generation of double or triple mutants using the traditional cross-breeding method. To elucidate the function of each LYK in nodulation, we designed five guide RNAs (gRNAs) based on conserved DNA regions across the three LYK genes and employed CRISPR/Cas technology to knock out these genes (Fig. 1C). Following multiple rounds of stable transformation assays, we successfully obtained all single, double, and triple mutants for the three genes (Fig. 1C and Fig. S1 online). Analysis of DNA insertions or deletions in each mutant line revealed frame-shift mutations leading to premature terminations of protein translation. Consequently, all these single, double, and triple mutant plants represent loss-of-function mutations for their respective genes.

To investigate the function of three LYKs during nodulation, we initially assessed nodulation phenotypes in all single mutants: *lyk2-1, lyk2-2, lyk3-1, lyk3-2, lyk2bis-1*, and *lyk2bis-2*, 21 days post-inoculation (dpi) with *Sinorhizobium meliloti* 2011. All single mutant plants exhibited similar nodule numbers compared to the WT plants (Fig. 1D). This observation aligns with previous reports indicating no discernible differences in nodule numbers between *Tnt1*-insertional mutants for each gene and WT plants [9]. Subsequently, we delved into nodulation phenotypes in three double mutant plants. Both the *lyk2/lyk2bis* and *lyk3/lyk2bis* double mutants failed to produce any nodules at 21 dpi, contrasting with the *lyk2/lyk3* mutant, which formed nodules comparable to WT plants (Fig. 1E). Further analysis of the triple mutant *lyk2/lyk3/lyk2bis* revealed an inability to form any nodules (Fig. 1F).

To gain insights into the inner cell structures of nodules generated on the *lyk2/lyk3* double mutant plants, we conducted toluidine blue staining assays for nodules. As depicted in Fig. 2E, the inner cell structures exhibited no observable differences in symbiotic cells between WT and *lyk2/lyk3* double mutant nodules (Fig. 2E). These findings collectively underscore the potential major role of MtLYK2bis in nodulation.

**Fig. 2.**
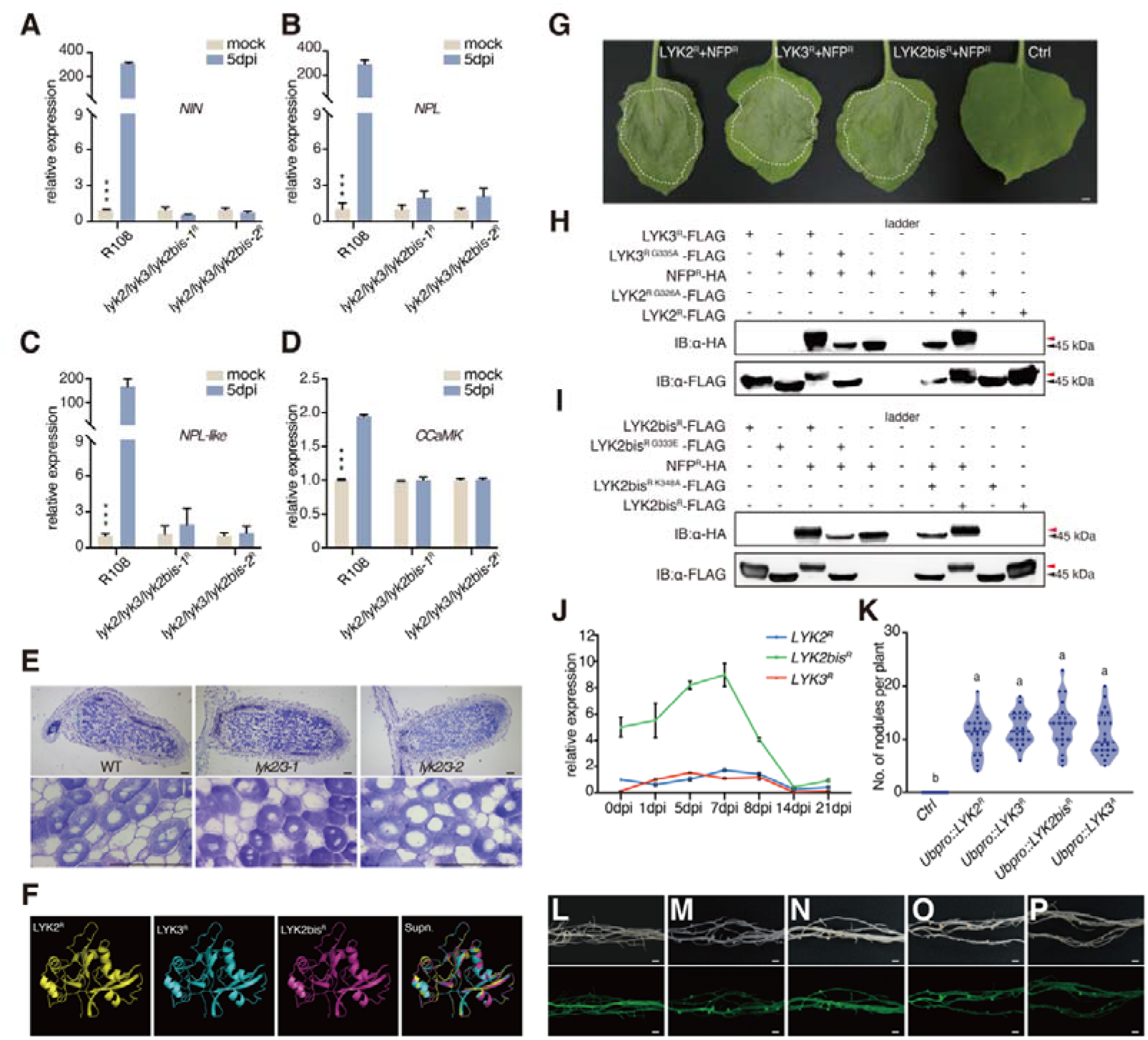
In *M. truncatula* R108, MtLYK2, MtLYK3, and MtLYK2bis possess similar physically and biochemical functions. (A, B, C and D) Quantification of expression of symbiosis-related genes in WT *M. truncatula* (R108) and two triple mutants upon *S. meliloti* 2011 inoculation using qPCR. In *M. truncatula* R108, the expression of *NIN* (A), *NPL* (B), *NPL-like* (C), and *CCaMK* (D) is upregulated in WT nodules 5 dpi with *Sinorhizobium meliloti* 2011 compared to triple mutant plants and uninoculated roots. (P < 0.05, one-way ANOVA test) Error bars represent SE (n = 3). (E) Cell structures of WT (R108) and the *lyk2/lyk3-1* and *lyk2/lyk3-2* mutant nodules. Toluidine blue-stained transverse sections of transgenic nodules at 21 dpi. From left to right, WT, *lyk2/lyk3-1, lyk2/lyk3-2* (Scale bars, 100μm). (F) Homology Models of R108-LYK2^ED^, R108-LYK3^ED^, and R108-LYK2bis ^ED^. These models were generated based on the crystal structure of A17-LYK3 ^ED^ (Protein Data Bank ID: 6XWE). From left to right: the individual models of R108-LYK2 ^ED^, R108-LYK3 ^ED^, R108-LYK2bis ^ED^, followed by a superposition of all three models. (G) MtLYK2, MtLYK3, and MtLYK2bis can physically interacts with NFP leads to cell death in *N. benthamiana* leaves at 3 d post-infiltration. (Scale bar, 3 mm) (H and I) MtLYK2, MtLYK3 and MtLYK2bis exhibit active kinase activity, inducing NFP phosphorylation and resulting in a constitutive band retardation. Phospho-deficient mutations lack phosphorylation activity and are unable to phosphorylate NFP. Red Arrowhead indicates band retardation. (J) Relative expression levels of *MtLYK2, MtLYK3*, and *MtLYK2bis* in nodules at various stages post-inoculation, determined using the comparative Ct method, relative to the expression of *MtLYK2* in un-inoculated conditions. Error bars represent SE (n = 3). (K) Complementation of nodulation capacity through the overexpression of MtLYK2, MtLYK3, MtLYK2bis, and A17-LYK3 in *hcl-1* mutants. Recording the nodulation phenotype in transformed roots. Analysis was conducted on 20 plants for each genotype/inoculum. Lowercase letters indicate significant differences (ANOVA, Tukey, P < 0.05). (L, M, N, O and P) Root transformation was monitored by the expression of a GFP marker gene and nodulation phenotype in bright (upper panel) and GFP-field (lower panel). (L): Ctrl (control), (M): *Ubpro::LYK2*, (N): *Ubpro::LYK3*, (O): *Ubpro::LYK2bis*, (P): *Ubpro::*A17*LYK3*. (Scale bar, 2 mm).

To comprehensively assess nodulation phenotypes, we examined physiological responses, including root hair deformation and infection thread formation, at the early stage of rhizobial infection in *M. truncatula* R108. In the *lyk2/lyk2bis* and *lyk3/lyk2bis* double mutant plants, 4 days post-inoculation (dpi) with GFP-labeled *S. meliloti* 2011, we observed root hair deformation and the presence of infection foci, but not the formation of infection threads (Fig. 1I and J), as compared to WT plants (Fig. 1H). However, in the *lyk2/lyk3/lyk2bis* triple mutant plants, we could not observe any root hair deformation, infection foci, or infection threads (Fig. 1K). Furthermore, we observed a complete block in the transcriptional induction of symbiosis-related genes, including *NIN, NPL, NPL-like*, and *CCaMK*, in the *lyk2/lyk3/lyk2bis* triple mutants compared to the high expression levels in WT plants (Fig. 2A, B, C, D). These data indicate that the *lyk2/lyk3/lyk2bis* triple mutants completely lose the response to rhizobium, mirroring observations in *L. japonicus nfr1* mutants. While both double mutant plants, *lyk3/lyk2bis* and *lyk2/lyk2bis*, failed to produce nodules, they retained responses to rhizobium attachment, albeit without establishing effective rhizobial invasion. These findings suggest that LYK2 and LYK3 have redundant roles in mediating rhizobium recognition and/or attachment.

To investigate the potential involvement of MtLYK2, MtLYK3, and MtLYK2bis in symbiotic signaling, we predict the protein structures of these receptors using the known structure of MtLYK3^ED^ (extracellular domain) as a template in SwissModel (Fig. 2F). The results revealed a high degree of similarity in three-dimensional structures among the three receptors. Previous reports have confirmed MtLYK2bis, which extends the nodulation specificity of R108 to the *Sinorhizobium meliloti nodF/nodL* mutant strain [9]. We separately inoculated three double mutants with *nodF/nodL* mutant strain and found that only *lyk2/lyk3-1* and *lyk2/lyk3-2* could generate a similar number of nodules as R108 (Fig. 1G). This indicates that MtLYK2bis can simultaneously function as both an entry receptor and a signaling receptor for recognizing *nodF/nodL* mutant strain.

The above data suggest that all three receptors, MtLYK2, MtLYK3, and MtLYK2bis, may possess redundant functions in mediating symbiotic signaling transduction. An intriguing observation arises from the co-expression of NFR1/MtLYK3 and NFR5/MtNFP (NF perception in *M. truncatula*, the ortholog of NFR5), which induces cell death in *Nicotiana benthamiana* leaves [3, 10]. To probe the associations of these three receptors with NFP, we co-overexpressed MtNFP with either MtLYK2, MtLYK3, or MtLYK2bis in *N. benthamiana*. Remarkably, we observed apparent cell death in leaves upon co-overexpressing MtNFP with any of MtLYK2, MtLYK3, or MtLYK2bis, compared to the control (Fig. 2G).

Given the established transphosphorylation between NFR1/MtLYK3 and NFR5/MtNFP in transducing NF-signaling in plant cells [3, 11], we investigated whether MtLYK2, MtLYK3, and MtLYK2bis share similar functions in phosphorylating MtNFP. Co-expression experiments involving the cytoplasmic domains (CDs) of MtLYK2, MtLYK3, and MtLYK2bis, along with their kinase-dead versions, and the CD of NFP or controls were conducted in Escherichia coli cells. The results revealed an apparent band retardation of NFP^CD^ in the presence of either MtLYK3^CD^, MtLYK2^CD^, or MtLYK2bis^CD^, but not their kinase-dead versions (Fig. 2H and I). Immunoblots further also identified the retardation of bands for MtLYK2^CD^, MtLYK3^CD^, and MtLYK2bis^CD^, indicating their capacity for kinase activity with auto-transphosphorylation, as well as trans-phosphorylation of MtNFP.

The above data suggest that MtLYK2, MtLYK3, and MtLYK2bis exhibit redundant functions, with MtLYK2bis playing a major role in mediating nodulation in *M. truncatula*. This observation prompted us to speculate about the differential regulation of transcription levels for *MtLYK2, MtLYK3*, and *MtLYK2bis*. To explore this speculation, we examined the transcriptional levels of *MtLYK2, MtLYK3*, and *MtLYK2bis* in *M. truncatula* roots and nodules at different time points post-inoculation with rhizobium. Both *MtLYK2* and *MtLYK3* displayed low expression levels in the roots before and after rhizobial treatment. In contrast, the expression of *MtLYK2bis* was significantly higher than that of *MtLYK2* and *MtLYK3* (Fig. 2J). The elevated expression levels of *MtLYK2bis* may contribute to its prominent role in mediating nodulation in *M. truncatula*.

To validate this hypothesis that all three receptors have redundant functions, we conducted complementation assays by expressing *MtLYK2, MtLYK3*, and *MtLYK2bis* driven by ubiquitin promoter in the *lyk2/lyk3/lyk2bis-1* triple mutant plants. Intriguingly, the overexpression of any of *MtLYK2, MtLYK3*, or *MtLYK2bis* completely restored nodulation in the *lyk2/lyk3/lyk2bis* mutant plants. Similar nodules were formed on each transgenic root compared with control plants (Fig. 2K). These data strongly suggest that all three LysM receptors share a similar function in mediating nodulation in *M. truncatula*.

In conclusion, our comprehensive study on multiple *M. truncatula* mutant plants has revealed that all three receptors, MtLYK2, MtLYK3, and MtLYK2bis, exhibit redundant functions in mediating nodulation. The overexpression of any of these receptors could complement nodulation defects in the *lyk2/lyk3/lyk2bis* mutant plants. Notably, among these receptors, MtLYK2bis appears to play a major role in nodulation, likely due to its relatively high expression levels compared with the other two genes in response to rhizobial inoculation. The differential expression patterns of the three LYK genes may be crucial for the precise regulation of nodulation in *M. truncatula*, although the underlying mechanisms are yet to be determined. Therefore, a one-receptor model with multiple specificities that mediates both signaling and entry responses is more suitable in *M. truncatula*.

## Acknowledgments

This work was supported by the National Key R&D Program of China (2019YFA0904700), the National Natural Science Foundation of China (32090063), and a Self-Innovation grant from the National Laboratory (AML2023B01). The qPCR and microscopy data are acquired from the Core Facility Center run by the National Lab of Agricultural Microbiology.

## Author contributions

Y.C. and Y.L. conceived the project. Y.L., Y.Z. and Z.Y. performed all the experiments. R.D. and H.Y. provided substantial help in experiments. H.Y. and H.Z. provided valuable suggestions for the study. Y.L., H.Z., and Y.C. analyzed the data and finalized all figures with inputs and comments from co-authors. Y.L. and Y.C. wrote the article.

