## Supplementary material for "Nodulation Trio in *Medicago truncatula:* Unveiling the Overlapping Roles of MtLYK2, MtLYK3, and MtLYK2bis": Fig. S1 online

**This PDF file includes:**

Supplementary Information, Materials and Methods
SI References
Fig. S1
Datasets S1

**Supplementary Information, Materials and Methods**

**Genome Collinearity Analysis and Phylogenetic Reconstruction.** Genome collinearity analysis performed between Medicago truncatula strains A17 and R108, utilizing MCScanX for the analysis and JCVI for visualization. The orthogroups comprising LYK-I receptor kinases in seven species, as identified in a previous study [1], were included. The R108 protein sequence, derived from unpublished data. A complete list of species and accession numbers can be found in Dataset S1. These sequences were aligned using ClustalW in MEGAX software [2], and a phylogenetic tree was constructed using the Neighbor-Joining (NJ) method with 1000 bootstrap replications. The resulting tree was visualized and refined using the iTOL online platform (https://itol.embl.de/tree/).

**Plant Material, Bacterial Strains, and Growth Conditions.** Plant growth conditions and rhizobia inoculation were conducted in accordance with established protocols [3]. The rhizobia strains used, including *S. meliloti* 2011, *S. meliloti* 2011-GFP, and *S. meliloti* 2011 *nodF/nodL*, were selected for inoculation experiments.

**Plasmid Construction.** For gene knockout experiments, the CRISPR/Cas9 vector was constructed following a previously reported method [3]. Briefly, the ubiquitin promoter was amplified from the pUB-GFP-3xFLAG vector and seamlessly integrated into the KpnI/XhoI sites of the pBluescript sk(+)−2 × 35S–Cas9 vector, replacing the 2 × 35S promoter. LjUbipro-Cas9 was then introduced into the *Kpn*I/*Xba*I sites of the pCAMBIA1302-sGFP. Subsequently, LjU6-gRNA fragments from the pBluescript SK(+)−LjU6-tRNA–gRNA vector were cloned into this construct. This assembly was utilized for stable transformation in Medicago, with CRISPR/Cas9 targets designed using CRISPR-P 2.0 (http://crispr.hzau.edu.cn/CRISPR2/). Relevant primer sequences are provided below:

CRISPR-gRNA1/2-F:

5'-TCCCGGCTGGTGCAGTCTATTATGCAGAACTTAGGTTTTAGAGCTAGAA-3'

CRISPR-gRNA1/2-R:
5'-TTCTAGCTCTAAAACTAATCTTTTGAAATCTGGCTTGCACCAGCCGGGA-3'

CRISPR-gRNA3/4-F:

5'-TCCCGGCTGGTGCAACTACCAGTAGCATGCCCACGTTTTAGAGCTAGAA-3'

CRISPR-gRNA3/4-R:

5'-TTGTCAATATGTTCATATACTGCACCAGCCGGG-3'

CRISPR-gRNA4/5-F:

5'-GTATATGAACATATTGACAAGTTTTAGAGCTAGAA-3'

CRISPR-gRNA4/5-R:

5'-TTCTAGCTCTAAAACTTACCATGGTCTAGTAGAGTTGCACCAGCCGGGA-3'

***Medicago truncatula* Mutant Plants.** Stable Medicago truncatula transgenic lines were developed using leaf explants, which were wounded and infected with *Agrobacterium tumefaciens* EHA105. Mutants were identified by sequencing targeted genes in primary transformants. Hygromycin selection was utilized as part of the procedure [3]. gRNA editing efficacy was verified using the following primers:

CR-LYK2gRNA1-F:	5'-AACATTCCATTCCCATGTGAATG-3'

CR-LYK2gRNA1-R:	5'-GATCACAACATCAAAAGGTGATG-3'

CR-LYK2gRNA2-F:	5'-CACACATGTATCAATATTTGCGAGAG-3'

CR-LYK2gRNA2-R:	5'-TCTCCACAAAATCACACGGGA-3'

CR-LYK2gRNA4/5-F:	5'-ACAATTCAAGTTCTTGTCAAAGTCA-3'

CR-LYK2gRNA4/5-R:	5'-TGGTCAAGCCAAAATCTGCA-3'

CR-LYK2bisgRNA1-F:	5'-AACATTCCATTCCCATGTGAATG-3'

CR-LYK2bisgRNA1-R:	5'-CCTTTGCTGAAATTGGCATCTAG-3'

CR-LYK2bisgRNA2/3-F:	5'-TCAGTATTTGAGAGAGCTTGCGAA-3'

CR-LYK2bisgRNA2/3-R:	5'-TCGCTAACTGTGTAGGACAGA-3'

CR-LYK2bisgRNA4/5-F:	5'-AATAATTTGAAACACACTGCAGAAC-3'

CR-LYK2bisgRNA4/5-R:	5'-AGTGACATATGTATTGTGAGATGTG-3'

CR-LYK3gRNA1-F:	5'-GGCGTAGTATTCGATAAATCTGG-3'

CR-LYK3gRNA1-R:	5'-CAAGTTGATTGGAAGTACCAAGTTC-3'

CR-LYK3gRNA2-F:	5'-GTATCAGTATCTGAGAGTGCTCAC-3'

CR-LYK3gRNA2-R:	5'-ACAACTCTGGTTATTTGCTTTGC-3'

CR-LYK3gRNA5-F:	5'-GCAAAGCAAATAACCAGAGTTGTG-3'

CR-LYK3gRNA5-R:	5'-ACCAATCATAGTTCAGTTGATCTCG-3'

***Medicago truncatula* Nodulation Assays.** Pot-based nodulation assays with *S. meliloti* 2011 were conducted over a 21-day period. Toluidine blue staining of Medicago nodule sections, using 20 to 30 samples, was performed as previously described [4].

**Fluorescence Microscopy.** For imaging the fluorescence of *S. meliloti* 2011-GFP, an Olympus FV1000 confocal laser-scanning microscope was used, with an excitation wavelength of 488 nm.

**Gene Expression Analysis.** Gene expression analysis in R108 and lyk2/lyk3/lyk2bis mutant plants was performed post-rhizobia treatment at various time points. Total RNA from roots was isolated using the RNeasy Plant Mini Kit (Yeasen), and reverse transcribed into cDNA using the Rever Tra Ace qPCR RT Master Mix with gDNA remover (Yeasen), following the manufacturer's instructions. q-PCR was performed using the following primers:

qpcr-LYK2-F:	5'-GGGTGCAACTGTGGTTATAATAATAGCA-3'

qpcr-LYK2-R:	5'-CATCTTGAGTTGAAAGCGCTTCAG-3'

qpcr-LYK2bis-F:	5'-CGAAAATTGCGACCAAAGCTG-3'

qpcr-LYK2bis-R:	5'-CTGCCATAGCTATACCAACAGC-3'

qpcr-LYK3-F:	5'-CGATACTCTTCAGAAGATTGCAAACC-3'

qpcr-LYK3-R:	5'-CTGCCATAGCTATACCAACAGC-3'

qpcr-NIN-F:	5'-ACCCTCATCAATTACCGTCCAC-3'

qpcr-NIN-R:	5'-GTTCAGCGCTGCTTCATATAGC-3'

qpcr-NPL-F:	5'-CAATCGGTGGAAGTTCGTCC-3'

qpcr-NPL-R:	5'-TCCAATTCCAGTTCTCCCACTC-3'

qpcr-NPLlike-F:	5'-ACAACCGCAAGAGACTAGCA-3'

qpcr-NPLlike-R:	5'-CCTGGCTTAGGATTCACAGGG-3'

qpcr-CCaMK-F:	5'-ACCGTTGATCCTAGCAAGAGAC-3'

qpcr-CCaMK-R:	5'-AGCTGCACGAAGTTTACGTC-3'

qpcr-Histone2A-F:	5'-TGCTCAGGACTTCAAGACTGAC-3'

qpcr-Histone2A-R:	5'-CTTCAGCAGCCTCTTGCAAT-3'

**Structural Modeling.** Homology modeling was performed with SWISS-MODEL using the crystal structure of A17LYK3^ED^(Protein Data Bank ID: 6XWE) as template, and visualized in PyMOL, Version 2.5.2(https://swissmodel.expasy.org/).

**Agroinfiltration on *Nicotiana benthamiana* Leaves.** *Agrobacterium tumefaciens* EHA105 containing ProCVMV:LYK2-3xFLAG, ProCVMV:LYK3-3xFLAG, ProCVMV:LYK2bis-3xFLAG, ProCVMV:LYK2bis-3xFLAG and ProCVMV:NFP-3xHA fusion constructs were used to agroinfiltrate the four oldest leaves of each *N. benthamiana* plant.

**Hairy root transformation.** Hairy root transformation was conducted by a standard procedure (Medicago Handbook).Using *A. rhizogenes*-mediated transformation , seedlings of *lyk2/3/2bis-1* were transformed using strains containing empty vector (Ctrl), ProCVMV:LYK2-3xFLAG, ProCVMV:LYK3-3xFLAG, ProCVMV:LYK2bis-3xFLAG and ProCVMV:LYK3-A17-3xFLAG constructs, transformants were selected by GFP. GFP-positive roots were selected and all the GFP-negative roots were removed. The nodulation phenotype was examined 4 wk after inoculation.

**Kinase Activity Assays.** Kinase activity was assessed via in vitro phosphorylation assays in E. coli. LYK2^CD^,LYK3^CD^,LYK2bis^CD^, and their point mutation variants were cloned into pET-28a(+) to generate C-terminal FLAG-tagged proteins. NFP^CD^ was tagged with HA and inserted into pCDFDuet-1. These constructs were co-transformed or separately transformed into *E. coli* BL21-CodonPlus RIL, and protein expression was induced with 0.3 mmol/L isopropyl β-D-1-thiogalactopyranoside at 18 ℃ for 8 hours.

[1]	Buendia L, Girardin A, Wang T, et al. Lysm receptor-like kinase and lysm receptor-like protein families: An update on phylogeny and functional characterization. Front Plant Sci, 2018, 9: 1531.

[2]	Tamura K, Stecher G, Kumar S. Mega11: Molecular evolutionary genetics analysis version 11. Mol Biol Evol, 2021, 38: 3022-3027.

[3]	Yu H, Xiao A, Dong R, et al. Suppression of innate immunity mediated by the cdpk-rboh complex is required for rhizobial colonization in medicago truncatula nodules. New Phytol, 2018, 220: 425-434.

[4]	Wang C, Yu H, Luo L, et al. Nodules with activated defense 1 is required for maintenance of rhizobial endosymbiosis in medicago truncatula. New Phytol, 2016, 212: 176-191.

**Fig. S1.** List of mutatant lines generated in this study. Deleted nuceotides are marked in red color, while inserted nucleotides are marked in blue color.

Deletion and/or insetion nuceotides	Location of the mutant sites

| Single mutant | *lyk2*-1 | 5'-GCTGTCTATTATGCAGAACTTAGAGGCGAGGT-3' | 5bp deletion in the 5th exon |
| --- | --- | --- | --- |
|  | *lyk2*-2 | 5'-TCTATTATGCAGAACTTAGAGGCGAGGTACG-3' | 1bp deletion in the 5th exon |
|  | *lyk3*-1 | 5'-TGTCTATTATGCAGAACTTAGAGGCGAGGTA-3' | 1bp deletion in the 4th exon |
|  | *lyk3*-2 | 5'-CAGGATCCAGTGCGGCATGCTACAGGTAGT-3' | 1bp insertion in the 4th exon |
|  | *lyk2bis* -1 | 5'-AGGTTCCAGTGGGCATGCTACTGGTAGTG-3' | 1bp deletion in the 5th exon |
|  | *lyk2bis* -2 | 5'-TCAGGTTCCAGTGTGGCATGCTACTGGTAT-3' | 1bp insertion in the 5th exon |
| Double mutant | *lyk2*  */lyk3*-1 | 5'-CTATTATGCAGAACTTAGAGGCGAGGTAC-3' | 1bp deletion in the 5th exon |
|  |  | 5'-AGCCAGATTTCAAAAGATTATGGCTTGTTT-3' | 1bp deletion in the 1st exon |
|  | *lyk2*  */lyk3*-2 | 5'-GAACTAGCAAA·······90bp········CTACATGAAT-3' | 90bp deletion flanking 5th exon and intron |
|  |  | 5'-AGCTGTCTATTATGCAGAACTTAGAGGCGA-3' | 1bp deletion in the 5th exon |
|  | *lyk2 /lyk2bis* -1 | 5'-CTGTCTATTATGCAGAACTTAGAGGCGAGG-3' | 5bp deletion in the 5th exon |
|  |  | 5'-CAAAAGAGGTATGTTATGGCTTGTTTGTTACCT-3' | 19bp deletion & 6bp insertion in the 5th exon |
|  | *lyk3 /lyk2bis* -1 | 5'-GGTACAGAACCATTACCATGGTCTAGTAGA-3' | 1bp deletion in the 8th exon |
|  |  | 5'-ACTTCAGGTTCCAGTGGGCATGCTACTGGTA-3' | 1bp deletion in the 5th exon |
|  | *lyk3 /lyk2bis* -2 | 5'-TACAGAACCATTACCATGGTCTAGTAGAGTG-3' | 2bp deletion in the 8th exon |
|  |  | 5'-GTACAGAACCATTACCATGGTCTAGTAGAGTG-3' | 8bp deletion in the 8th exon |
| Triple mutant | *lyk2 /lyk3*  */lyk2bis* -1 | 5'-TATTATGCAGAACTTAGAGGCGAGGTACGAAA-3' | 19bp deletion in the 5th exon |
|  |  | 5'-GCTGTCTATTATGCAGAACTTAGAGGCGAGGT-3' | 1bp deletion in the 1st exon |
|  |  | 5'-GCTGTCTATTATGCAGAACTTAGAGGCGAGGT-3' | 7bp deletion in the 5th exon |
|  | *lyk2 /lyk3*  */lyk2bis* -2 | 5'-GCTGTCTATTATGCAGAACTTAGAGGCGAGGT-3' | 1bp deletion in the 5th exon |
|  |  | 5'-GAAAATAGCTT·······592bp········TGTGAGTTGA-3' | 592bp deletion flanking 5th exon and two introns |
|  |  | 5'-CTGTCTATTATGCAGAACTTAGAGGCGAGGTA-3' | 7bp deletion in the 5th exon |

**Datasets S1** All the LYK sequences used for for phylogenetic tree analysis in Fig. 1B.

>Solanum lycopersicum LYK13

-----------------------MIFLRRRSITIL----VLIYFF----SNCTTCYSTSCTNGCDLALASFF--IWPE-SNLPLINQLFDNISY-------SDILEW-NTQ--ITSTFILTESRVHVPFRCDCLN-----NGEFLGHVFSYNVSA----NETYDLIATRRYSSLTNKELLM--RDNRYPDNNIP-DHVTLNVTVNCSCGNKHVSKDYGLFITYPMRPGENLSYIALVTNTSS----KLIEMYNPMVN--FSAGSG-LLYIPGRDKLGNYPPIST--RK-G-------------------------------------------SSGKTIAALAVASLAGVLLLVGIIYVGIY--RRKEQKVAANIPVSSGQCY---PPSP-----------------------GLSGIHVDKSVEFSYQELAESTDNFSISNKIGEGGFGAVYYAELRGKKAAIKRMNREGRTEFLAELKILTRVHHLNLVSLIGYCVERSLFLVYEFIENGNLSQHLHG-----RDVLTWSTRVQIAMDSARGLEYIHEHTVPFYIHRDVKSANILINKNFHAKIGDFGLSKLVESGNPTLNT--RFMGTFGYMPPEYGHSGVI--SRKVDVYAFGVVLYELISSKDAI--------------VKEDG--VDEARSLVALFDEAHSHP-NQIEAISRLIDPKLCDDYPLDSVYKMAQLAKSCTEKNPEMRPTMKSVVVALMALSSS-HA--------------------------------

>Arabidopsis thaliana CERK1

---------------------------MKLKISLIAPILLLFSFF-------FAVESK-CRTSCPLALASYY--LENG-TTLSVINQNLNSSIAPYDQINFDPILRY-NSN-IKDKDRIQMGSRVLVPFPCECQ------PGDFLGHNFSYSVRQ----EDTYERVAISNYANLTTMESLQ--ARNPFPATNIP-LSATLNVLVNCSCGDESVSKDFGLFVTYPLRPEDSLSSIARSSGVSA----DILQRYNPGVN--FNSGNG-IVYVPGRDPNGAFPPFKS-SKQDG------------------------------------------VGAG-VIAGIVIGVIVALLLILFIVYYAY---RKNKS-KGDSFSSS--------IPLSTKADHASSTSLQSGGLGGAGVSPGIAAISVDKSVEFSLEELAKATDNFNLSFKIGQGGFGAVYYAELRGEKAAIKKMDMEASKQFLAELKVLTRVHHVNLVRLIGYCVEGSLFLVYEYVENGNLGQHLHG--SG-REPLPWTKRVQIALDSARGLEYIHEHTVPVYVHRDIKSANILIDQKFRAKVADFGLTKLTEVGGSATR---GAMGTFGYMAPETV-YGEV--SAKVDVYAFGVVLYELISAKGAV--------------VKMTE-AVGEFRGLVGVFEESFKET-DKEEALRKIIDPRLGDSYPFDSVYKMAELGKACTQENAQLRPSMRYIVVALSTLFSS-TG--NWDVGNF-QNEDLVSLMSGR----------

>Oryza sativa LYK10

------------------------------MFSLP----ALLIGACAFAAAAVAASGDGCRAGCSLAIAAYY--FSEG-SNLTFIATIFAIGGGGY-----QALLPY-NPA-ITNPDYVVTGDRVLVPFPCSCLGLPAAPASTFLAGAIPYPLPLPRGGGDTYDAVAA-NYADLTTAAWLE--ATNAYPPGRIPGGDGRVNVTINCSCGDERVSPRYGLFLTYPLWDGETLESVAAQYGFSSPAEMELIRRYNPGMG--GVSGKG-IVFIPVKDPNGSYHPLKS--GGMGNS----------------------------------------LSGG-AIAGIVIACIA-IFIVAIWLIIMFY--RWQKFRKATSRPSPEE------TSHLDDAS-------------------QAEGIKVERSIEFSYEEIFNATQGFSMEHKIGQGGFGSVYYAELRGEKTAIKKMGMQATQEFLAELKVLTHVHHLNLVRLIGYCVENCLFLVYEFIDNGNLSQHLQR--TG-YAPLSWATRVQIALDSARGLEYLHEHVVPVYVHRDIKSANILLDKDFRAKIADFGLAKLTEVGSMSQSLSTRVAGTFGYMPPE-ARYGEV--SPKVDVYAFGVVLYELLSAKQAI--------------VRSSE-SVSESKGLVFLFEEALSAP-NPTEALDELIDPSLQGDYPVDSALKIASLAKSCTHEEPGMRPTMRSVVVALMALTAN-TDLRDMDYHPF-----------------------

>Oryza sativa LYK9

--------------------------MEASTSLLV----LVLAAAAFAAGTVTEAAGDGCSAGCDLALASFY--VTPN-QNVTNMADLFGIGAANY-----RSLAPY-NPN-IPNLDFINVGGRVNVYFTCGCRSLPGSPGATYLAGAFPFQMSR----GQIYTSVAA-NYNNLTTAEWLQ--ATNSYPANNIP-DTAVINATVNCSCGDASISPDYGLFLTYPLRAEDTLASVAATYGLSS--QLDVVRRYNPGME--SATGSG-IVYIPVKDPNGSYLPLKS--PGKG------------------------------------------ASAG-AIAGGVVAGV--VVLAAIFLYIIFY--RRRKA-KQATLLQSSED-----STQLGTISMDKVTPSTIVGPS------PVAGITVDKSVEFSYEELSNATQGFSIGNKIGQGGFGAVYYAELRGEKAAIKKMDMQATHEFLAELKVLTHVHHLNLVRLIGYCIESSLFLVYEFIENGNLSQHLRG--MG-YEPLSWAARIQIALDSARGLEYIHEHTVPVYIHRDIKSANILIDKNYRAKVADFGLTKLTEVGGTSMPTGTRVVGTFGYMPPEYARYGDV--SPKVDVYAFGVVLYELISAKEAI--------------VRSTE-SSSDSKGLVYLFEEALNSP-DPKEGLRTLIDPKLGEDYPIDSILKLTQLAKVCTQEDPKLRPSMRSVVVALMTLSST-SE--FWDMNNLYENQGLVNLMSGR----------

>Solanum lycopersicum LYK11

----------------------MIFFQENRSFELV----LGLLVL---NILWVGVKSQ-CSDDCD-ALASFY--VWNG-ANLTFMSNTFSTPI--------KNILSY-NPQ-ITNPDIIQSQSRVNVPFSCSCV------DGKFMGHQFDVQVKT----NTTYPRITRLYCSNLTTVEKLQ--ESNSYDPNNVP-VNSIVKVIVNCSCGNSHVSKDYGLFITYPLRPGENLVTLANDFSLPQ----KLLEDYNPEAN--FSSGSG-LVFIPGKDQNGTYPQLRTSTSSKG------------------------------------------FSGG-AITGISVAVVLVVALLAVCIYITFY--RGRKTEENLNLEPYKH------SSNKHIPGHANFENSSEGGSLKQGASPEVPRIAVDKSIEFSYDELAKASDNFSTAYKIGQGGFASVYYGELRGEKAAIKKMDMQATKEFLAELKVLTHVHHLNLVRLIGYCVEGSLFLVYEYIENGNLSQHLRGFVPG-KVPLPWSTRVKIALDAARGLEYIHEHTVPVYIHRDIKTANILIDKNFRAKVADFGLTKLIETEGGSMNT--RLVGTFGYMAPEYGQFGNV--SLKIDVYAFGVVLYELISARKAI--------------IKTSE-ISTESKGLVGLFEDVLNEV-DPKEGICKLVDPKLGDDYPLDSVWNVALLAKACTQENPQLRPSMRSIVVALMTISSTSTA--DWNLGEFYENQGLAHLISGR----------

>Lupinus angustifolius LYK1

---------------------------MKLK--------LVFLLL--LKYVCFIVESK-CIKGCDLALASYYVPVWPI-VSLGNITSFMHSNVLTNP----NVVTSY-NKDKVFNGDVMLALYRTNVPFPCDCI------GGEFLGHVFEYSAVE----GDTYGLIAMKRYSNLTTVEILK--RFNSYDPNHIP-VNAKVNVTVKCSCGNSQISKDYGLFITYPLRPGNNLQELSKETKIDA----KLLQSYNPGVN--FSQENG-IVFIPGKDQNGVYVPLYP--RTGG------------------------------------------VAKG-VAVGISIAATCGLVLLVICIYDRYF--KKKEG-EKAKLSIENSIGF---STQD--AYGSGEYETSGSSVHAS----ALTGIMVAKSLEFSYQELAKATNNFSLDNKIGQGGFGAVYYAELRGEKTAIKKMDVQASSEFLAELKVLTHVHHLNLVRLIGYCVEGSLFLVYEYIDNGNLGQYLHG--KG-KDPLPWSTRLQIALDSARGLEYIHEHTVPLYIHRDVKSANILIDKNLRGKVADFGLTKLIEVGTSSLHT--RLVGTFGYMPPEYAQYGDI--SPKIDVYAFGVVLYELISAKNAV--------------LKTGE-TVAESKGLVNLFEEALNQI-NPLEPLTTLVDPRLGDNYPIQSLLKIAELGRACTRDNPLLRPNMKSIVVALMTLSSS-NE--D-NTTSSYDNQTLINLLLDEGFRGITF---

>Lupinus angustifolius LYK2

---------------------------MKLK--------FVFFLL--LECVCFIVESK-CIKGCDLALASYYVSVWPS-ISLGNITNFMHSNVLTNS----DVIISY-NKGKIFNGDVLLSLTRTNVPFPCDCI------GGEFLGHVFQYSSVA----GDTYDTIAMKSYSNLTTVEFLK--RFNSYDPNHIP-VNSKVNVTINCSCGNSLISKDYGLFTTYPLRPGNNLQELSKETNIDA----KLLQSYNPGAN--FSQESR-IVFIPGRDQNGVYVPLYP--RIGG------------------------------------------LARG-AAVGISIAATCGLVLLIICIYDRYF--KKKEG-EKTKLSTEDSIGL---STEDGTGSGSGEYEASGSSGHAV----GLTSIMVAKSLEFSYQELAKATNNFSLDNKIGQGGFGAVYYAELRGEKTAIKKMDVQASSEFLAELKVLTHIHHSNLVRLIGYCVEGSLFLVYEYIDNGNLGQYLHG--KG-RDPLPWSTRLQIALDSARGLEYIHEHTVPLYIHRDVKSANILIDKNLRGKVADFGLTKLIEVGTSSFHT--RLVGTFGYMPPEYAQYGDI--SSKIDVYAFGVVLYELISAKSAI--------------LKTGE-TVSESKGLVTLFEGALNQI-NPLEALPKLVDPRIGDNYPIESVLKIAQLGRACTRDNPLLRPNMKSIVVALMTLSSS-AE-----HSTSYDNQTLINLLLDEGFREIITPTS

>Mt A17 LYK2

---------------------------MKLKNGLL----LFF-MF--LECVFSKVESK-CVKGCDVALASYH--VMLP-FTYQNITSFMQSKIVSVSSLS-DVIISY-NKGKVSKNGNLFAFSRVNIPFPCECI------GGDFLGHVFEYSAKE----GDTYDLIANSYYASLTTVELLK--KFNSYDQDHIP-AKAKVNVTVNCSCGNSQISKDYGLFITYPLRTDDTLQKIANQSNLDE----GLIQSYNSGVN--FSNGSG-IVFIPGRDQNGDYVPLYP--RS-G------------------------------------------LAKG-ATVGIIIAGIFGLLLLVIYIYVRYF--KKKEE-EKTKL-----------AEALSTQDGSAEYETSGSSVHAT----VFTGIMVAKSTEFSYQELAKATNNFSLDNKIGQGGFGAVYYAELRGEKTAIKKMDVQASSEFLCELKVLTHVHHLNLVRLIGYCVEGSLFLVYEHIDNGNLGQYLHG--TG-KEPLPWSSRVEIALDSARGLEYIHEHTVPMYIHRDVKSANILIDKNLRGKVADFGLTKLLEVGNSTLQT--RLVGTFGYMPPEYAQYGDV--SPKIDVYAFGVVLFELISAKNAV--------------LKTGE-FVAESRGLVALFEEALNQT-DPLESLRKLVDPRLREDYPIDSVLKMAQLGRECTKDNPLLRPSMRSIVVSLMSLLSP-SE--DCDGDTSDENQTIINLLSVR----------

>Mt R108 LYK2

---------------------------MKLKNGLL----LFF-MF--LECVFSKVESK-CVKGCDVALASYH--VMLP-FTYQNITTFMQSKIVSVSSLS-DVIISY-NKDKVSKNGNLFAFSRVNIPFPCECI------GGEFLGHVFEYSAKE----GDTYDLIANSYYASLTTVELLK--KFNSYDQDHIP-AKAKVNVTVNCSCGNSQISKDYGLFITYPLRTDDTLQKIANQSNLDE----GLIQSYNSGVN--FSNGSG-IVFIPGRDQNGDYVPLYP--RS-G------------------------------------------LAKG-ATVVIIIAGIFGLLLLVIYIYVRYF--RKKEE-EKTKL-----------AEALSTQDGSAEYEISGSSVHAT----VFTGIMVAKSTEFSYQELAKATNNFSLDNKIGQGGFGAVYYAELRGEKTAIKKMDVQASSEFLCELKVLTHVHHLNLVRLIGYCVEGSLFLVYEHIDNGNLGQYLHG--TG-KEPLPWSSRVEIALDSARGLEYIHEHTVPMYIHRDVKSANILIDKNLRGKVADFGLTKLLEVGNSTLHT--RLVGTFGYMPPEYAQYGDV--SPKIDVYAFGVVLFELISAKNAV--------------LKTGE-FVAESRGLVVLFEEALNQT-DPLESLHKLVDPRLREDYPIDSVLKMAQLGRECTKDNPLLRPSMRSIVVSLMSLLSP-SE--DCDGDTSDENQTIINLLSVR----------

>Mt R108 LYK2bis

---------------------------MKLKNGLL----LFF-MF--LECVFSKVESK-CVKGCDVALASYH--VMLP-FTYQNITTFMQSKIVSVSSLS-DVIISY-NKDKVSKNGNLFAFSRVNIPFPCECI------GGEFLGHVFEYSAKE----GDTYDLIANSYYASLTTVELLK--KFNSYDQDHIP-AKAKVNVTVNCSCGNSQISKDYGLFVTYPLRSDDTLAKIATKADLDE----GLIQNFNLDAN--FSKGSG-IVFIPGRDQNGHYFPLYP--RT-G------------------------------------------IAKG-SAVGIAMAGIFGLLLFVIYIYAKYF--QKKEE-EKTKLPQTSR------AFSTQDASGSAEYETSGSSGHATGSAAGLTGIMVAKSTEFTYQELAKATNNFSLDNKIGQGGFGAVYYAELRGEKTAIKKMDVQASSEFLCELKVLTHVHHLNLVRLIGYCVEGSLFLVYEHIDNGNLGQYLHG--IG-TEPLPWSSRVQIALDSARGLEYIHEHTVPVYIHRDVKSANILIDKNLRGKVADFGLTKLIEVGNSTLHT--RLVGTFGYMPPEYAQYGDV--SPKIDVYAFGVVLYELITAKNAV--------------LKTDE-SVAESKGLVQLFEEALHQM-DPLEGLRKLVDPRLKENYPIDSVLKMAQLGRACTRDNPLLRPSMRSIVVALMTLSSP-TE--DCDDDSSYEKQSLINLLSTR----------

>Mt A17 LYK3

---------------------------MNLKNGL-----LLFILF--LDCVFFKVESK-CVKGCDVALASYY--IIPS-IQLRNISNFMQSKIVLTNSF--DVIMSY-NRDVVFDKSGLISYTRINVPFPCECI------GGEFLGHVFEYTTKE----GDDYDLIANTYYASLTTVELLK--KFNSYDPNHIP-VKAKINVTVICSCGNSQISKDYGLFVTYPLRSDDTLAKIATKAGLDE----GLIQNFNQDAN--FSIGSG-IVFIPGRDQNGHFFPLYS--RT-G------------------------------------------IAKG-SAVGIAMAGIFGLLLFVIYIYAKYF--QKKEE-EKTKLPQTSR------AFSTQDASGSAEYETSGSSGHATGSAAGLTGIMVAKSTEFTYQELAKATNNFSLDNKIGQGGFGAVYYAELRGEKTAIKKMDVQASSEFLCELKVLTHVHHLNLVRLIGYCVEGSLFLVYEHIDNGNLGQYLHG--IG-TEPLPWSSRVQIALDSARGLEYIHEHTVPVYIHRDVKSANILIDKNLRGKVADFGLTKLIEVGNSTLHT--RLVGTFGYMPPEYAQYGDV--SPKIDVYAFGVVLYELITAKNAV--------------LKTGE-SVAESKGLVQLFEEALHRM-DPLEGLRKLVDPRLKENYPIDSVLKMAQLGRACTRDNPLLRPSMRSIVVALMTLSSP-TE--DCDDDSSYENQSLINLLSTR----------

>Mt R108 LYK3

---------------------------MNLKNGLL----LFI-LF--LDCVFFKVESK-CVQECDVALASYY--VMRS-VQFRNVTNFMQSKIVLTNSF--DVIMSY-NRGVVFDKSGLISYTRVNVPFPCECI------GGEFLGHVFEYTAKE----GDDYDLIANTYYASLTTVELLK--KFNSYDPNHIP-VKAKINVTVNCSCGNSQISKDYGLFVTYPLRSDDTLQKIANQSNLDE----GLIQKYNSGAN--FSKGSG-IVFIPGRDQNGHYSYHIT-----G------------------------------------------IAKG-SAVGIAMAGIFGLLLFVIYIYAKYF--QKKEE-EKTKLPQTSR------AFSTQDASGSAEYETSGSSGHATGSAAGLTGIMVAKSTEFSYQELAKATNNFSLDNKIGQGGFGAVYYAELRGEKTAIKKMDVQASSEFLCELKVLTHVHHLNLVRLIGYCVEGSLFLVYEHIDNGNLGQYLHG--IG-TEPLPWSSRVQIALDSARGLEYIHEHTVPVYIHRDVKSANILIDKNLRGKVADFGLTKLIEVGNSTLHT--RLVGTFGYMPPEYAQYGDV--SPKIDVYAFGVVLYELITAKNAV--------------LKTGE-SVAESKGLVQLFEEALHQM-DPLEGLRKLVDPRLKENYPIDSVLKMAQLGRACTRDNPLLRPSMRSIVVALMTLSSP-TE--DCDDDSSYEKQSLINLLSTR----------

>Lotus japonicus NFR1

---------------------------MKLKTGLL----LFFILL--LGHVCFHVESN-CLKGCDLALASYY--ILPGVFILQNITTFMQSEIVSSN----DAITSY-NKDKILNDINIQSFQRLNIPFPCDCI------GGEFLGHVFEYSASK----GDTYETIANLYYANLTTVDLLK--RFNSYDPKNIP-VNAKVNVTVNCSCGNSQVSKDYGLFITYPIRPGDTLQDIANQSSLDA----GLIQSFNPSVN--FSKDSG-IAFIPGRYKNGVYVPLYH--RTAG------------------------------------------LASG-AAVGISIAGTFVLLLLAFCMYVRY---QKKEE-EKAKLPTDISMAL---STQDGNASSSAEYETSGSSGPGTASATGLTSIMVAKSMEFSYQELAKATNNFSLDNKIGQGGFGAVYYAELRGKKTAIKKMDVQASTEFLCELKVLTHVHHLNLVRLIGYCVEGSLFLVYEHIDNGNLGQYLHG--SG-KEPLPWSSRVQIALDAARGLEYIHEHTVPVYIHRDVKSANILIDKNLRGKVADFGLTKLIEVGNSTLQT--RLVGTFGYMPPEYAQYGDI--SPKIDVYAFGVVLFELISAKNAV--------------LKTGE-LVAESKGLVALFEEALNKS-DPCDALRKLVDPRLGENYPIDSVLKIAQLGRACTRDNPLLRPSMRSLVVALMTLSSL-TE--DCDDESSYESQTLINLLSVR----------

>Glycine max NFR1a

---------------------------MELKKGLL----VFF-LL--LECVCYNVESK-CVKGCDVAFASYY--VSPD-LSLENIARLMESSI--------EVIISF-NEDNISN-GYPLSFYRLNIPFPCDCI------GGEFLGHVFEYSASA----GDTYDSIAKVTYANLTTVELLR--RFNGYDQNGIP-ANARVNVTVNCSCGNSQVSKDYGMFITYPLRPGNNLHDIANEARLDA----QLLQRYNPGVN--FSKESG-TVFIPGRDQHGDYVPLYP--RKTG------------------------------------------LARG-AAVGISIAGICSFLLLVICLYGKYF--QKKEG-EKTKLPTENSMAF---STQD--VSGSAEYETSGSSGTA--SATGLTGIMVAKSMEFSYQELAKATNNFSLENKIGQGGFGAVYYAELRGEKTAIKKMDVQASTEFLCELKVLTHVHHFNLVRLIGYCVEGSLFLVYEYIDNGNLGQYLHG--TG-KDPLPWSGRVQIALDSARGLEYIHEHTVPVYIHRDVKSANILIDKNIRGKVADFGLTKLIEVGGSTLHT--RLVGTFGYMPPEYAQYGDI--SPKVDVYAFGVVLYELISAKNAV--------------LKTGE-SVAESKGLVALFEEALNQS-NPSESIRKLVDPRLGENYPIDSVLKIAQLGRACTRDNPLLRPSMRSIVVALMTLSSP-TE--DCD--TSYENQTLINLLSVR----------

>Glycine max NFR1b

---------------------------MELKKWL-----LFFLLL---EYVCCNAESK-CVKGCDVALASYY--VSPGYLLLENITRLMESIVLSN-----SDVIIY-NKDKIFN-ENVLAFSRLNIPFPCGCI------DGEFLGHVFEYSASA----GDTYDSIAKVTYANLTTVELLR--RFNSYDQNGIP-ANATVNVTVNCSCGNSQVSKDYGLFITYLLRPGNNLHDIANEARLDA----QLLQSYNPGVN--FSKESGDIVFIPGK----------------G------------------------------------------LATS-ASVGIPIAGIC-VLLLVICIYVKYF--QKKEG-EKAKLATENSMAF---STQD--VSGSAEYETSGSSGTASTSATGLTGIMVAKSMEFSYQELAKATNNFSLENKIGQGGFGIVYYAELRGEKTAIKKMDVQASTEFLCELKVLTHVHHLNLVRLIGYCVEGSLFLVYEYIDNGNLGQYLHG--TG-KDPFLWSSRVQIALDSARGLEYIHEHTVPVYIHRDVKSANILIDKNFRGKVADFGLTKLIEVGGSTLQT--RLVGTFGYMPPEYAQYGDI--SPKVDVYAFGVVLYELISAKNAV--------------LKTVE-SVAESKGLVALFEEALNQS-NPSESIRKLVDPRLGENYPIDSVLKIAQLGRACTRDNPLLRPSMRSIVVALLTLSSP-TE--DCYDDTSYENQTLINLLSVR----------

>Lotus japonicus LYS2

---------------------------MDLKSRLT----FFFLLS--WACISFSVVESMCISGCDLALASYY--IWIG-SNLTYISNIMESRVLSEP----EDIINY-NQDHVRNPDVLQVHTRVNVPFPCDCI------NGEFLGHIFLHEFHE----GDTYPSVAGTVFSNLTTDAWLQ--STNIYGPTSIP-VLAKVDVTVNCSCGDIKVSKDYGLFITYPLRAEDTLESIAEEAKLQP----HLLQRYNPGVD--FSRGNG-LVFIPGKDENGVYVPLHI--RKAG------------------------------------------LAR--VVAGVSIGGTCGLLLFALCIYMRYF--RKKEG-EEAKFPPKESMEP---SIQDDSKIHPAANGSA-----------GFKYIMMDRSSEFSYEELANATNDFNLANKIGQGGFGEVYYAELRGEKVAIKKMKIQASREFLAELKVLTSVHHLNLVRLIGYCVERSLFLVYEYMDNGNLSQHLRE--SE-RELMTWSTRLQIALDVARGLEYIHDYTVPVYIHRDIKPDNILLNKNFNAKVADFGLTKLTDIESSAINTD-HMAGTFGYMPPENA-LGRV--SRKIDVYAFGVVLYELISAKEAVVEIKESSTELKSLEIKTDE-PSVEFKSLVALFDEVIDHEGNPIEGLRKLVDPRLGENYSIDSIREMAQLAKACTDRDPKQRPPMRSVVVVLMALNSA-TD--DRMSHAEVNSSRAGALSPTVESL-------

>Mt A17 LYK1

--------------------------MKPIKFILS----LLLMLL-----ASSSAESK-CSKTCDLALASYY--IWEG-TNLTYISNIMQSNVVSKP----LDIFSY-NTDTLPNLDMLRFSSRLNVPFPCDCI------NDEFLGHTFLYEFHP----RETYASIAELTFSNLTNKEWME--K------VNVP-DSVKVNVTVNCSCGDKMVSKDYGLFITYPLSSEDTLESIAKHTKVKP----ELLQKYTPGVN--FSKGSG-LVFIPGKDKNGVYVPLPH--GKAGH-----------------------------------------LARS---LATAVGGTCTVLLLAISIYAIYF--RNKNA-KESKLPSK--------------------------------------YIVVDKSPKFSYEELANATDKFSLANKIGQGGFGEVYYGEPRGKKTAIKKMKMQATREFLAELKILTRVHHCNLVHLIGYCVEGSLFLVYEYIDNGNLSQNLHD--SE-RGPMTWSTRMQIALDVARGLEYIHEHSVPVYIHRDIKSDNILLNENFTGKIADFGLTRLTDSANSTDNTL-HVAGTFGYMPPENV-YGRI--SRKIDVYAFGVVLYELISAKPAVIKIDKTEFE---SEIRTNE-SIDEYKSLVALFDEVIDQKGDPIEGLRNLVDPRLEDNYSIDSISKMAKLARACLNRDPKRRPTMRAVVVSLMTLNST-ID--DGSRSASAALSTVMEHDSK-----------

>Mt R108 LYK1

--------------------------MKPIKFILS----LLLMLL-----ASSSAESK-CSKTCDLALASYY--IWEG-TNLTYISNIMQSNVVSKP----LDIFSY-NTDTLPNLDMLRVSSRLNVPFPCDCI------NDEFLGHTFLYEFHP----RETYASIAEFTFSNLTNKEWME--------KVNVP-DSVKVNVTVNCSCGDKMVSKDYGLFITYPLSSEDTLESIAKHTKVKP----ELLQKYNPGVN--FSKGSG-LVFIPGKDKNGVYVPLPH-----G------------------------------------------------SLATAVGGTCMVLLLAISIYAIYF--RNKNA-KETKLPSK--------------------------------------YIVVDKSPKFSYEELANATDNFSLANKIGQGGFGEVYYGEPRGKKTAIKKMKMQATREFLAELKILTRVHHCNLVHLIGYCVEGSLFLVYEYIDNGNLSQHLHD--SE-RGPMTWSTRMQIALDVARGLEYIHEHSVPVYIHRDIKSDNILLNENFTVKIADFGLTRLTDSANSTDNTL-HVAGTFGYMPPENV-YGRI--SRKIDVYAFGVVLYELISAKPAVFKIDKTEFG---SEIRTNE-SIDEYKSLVALFDEVIDQKGDPIEGLRNLVDPRLGDNYSIDSISKMAKLARACLNRDPKRRPTMRAVVVFLMTLNST-ID--DGSRTASAALSTVMEHDLN-----------

>Mt A17 LYK4

---------------------------MNLKNGL-----LLFILF--LDCVFFKVETK-CVKGCDVALASYY--IMPS-IQLINVSNFIQSKIVLTNSF--DVIMSY-NRVVVFDKSGLISYTRINVPFPCECI------GGEFLGHVFEYTTKE----GDDYDLIANTYYASLTTVELLK--KFNSYDPNHIP-VKAKINVTVICSCGNSQISKDFGLFVTYPLRSDDTLAKIATKADLDE----GLLQNFNQDAN--FSKGSG-IVFIPGRDENGVYVPLPS--RKAGH-----------------------------------------LARSLVAAGICIRGVCMVLLLAICIYVRYF--RKKNG-EESKLPPEDSMSP---STKDGDKDSYSDT--------------RSKYILVDKSPKFSYKVLANATENFSLAKKIGQGGFGEVYYGVLGGKKVAIKKMKTQATREFLSELKVLTSVRHLNLVHLIGYCVEGFLFLVYEYMENGNLSQHLHN--SE-KELMTLSRRMKIALDVARGLEYIHDHSVPVYIHRDIKSDNILLNKNFNGKIADFGLTKLTNIANSTDNTN-HMAGTFGYMPPENA-YGRI--SRKMDVYAFGVVLYELISAKAAVIMIDKNEFE--SHEIKTNE-STDEYKSLVALFDEVMDQKGDPIEGLRKLVDPRLGDNYSIDSISKMAKLAKACINRDPKQRPKMRDVVVSLMKLIST-ID--DESRTDSAELSLDVEHDSN-----------

>Mt R108 LYK5bis

MYIIRPNTLLHIYSTHPNSYSVCYTMKPIIKFRLS----LLF-LL--LVSKSITSESK-CNKTCDLALASYY--IRPG-TTLANISKVMQSNVVSKA----EDISIY-NIYTILSVDSVQVYTRLNVPFPCDCI------NDEFLGHTFLYKLRH----GDSYASIAKTTFGNLVTEEWIE--RVNVYSRTYVP-DSVMINVTVNCSCGNGEVSKDYGLFITYPLRPDDTLESIAKYTKVKG----ELLQRYNPGVN--FSQGRG-LVYIPGKDENGVYVPLPS--RKAGH-----------------------------------------LARSLVAAGICIRGVCMVLLLAICIYVRYF--RKKNG-EESKLPPEDSMSP---STKDGDKDSYSDT--------------RSKYILVDKSPKFSYKELANATENFSLAKKIGQGGFGEVYYGVLGGKKVAIKKMKMQATREFLSELKVLTSVYHLNLVNLIGYCVEGFLFLVYEYMENGNLSQHLHN--SE-KELMTLSRRMKIALDVARGLEYIHDHSVPVYIHRDIKSDNILLNKNFNGKIADFGLTKLTNIANSTDNTN-HMAGTFGYMPPENA-YGRI--SRKMDVYAFGVVLYELISSKAAVIMIDKNEFE--SHEIKTNE-SIDEYKSLVALFDEVMDQTGDPIEGLRKLVDPRLGDNYSIDSISKLAILAKACVNRDPKQRPKMRDVVVSLMKLNST-ID--DESRTGSAELSLAVEHDSN-----------

>Mt A17 LYK5bis

-------------------------MKPIIKFRLS----LLF-LL--LVSQSITSESK-CNKTCDLALASYY--IRPG-TTLANISKVMQSNVVSKA----EDISIY-NIYTILSVDSVQVYTRLNVPFPCDCI------NDEFLGHTFLYKLRH----GDSYASIATTTFGNLVTEEWIE--RVNVYPRTYVP-DSVMINVTVNCSCGNGEVSKDYGLFITYPLRPDDTLESIAKYTKVKG----ELLQRYNPGVN--FSQGRG-LVYIPGKDENGVYVPLPS--RKAGH-----------------------------------------LARSLVAAGICIRGVCMVLLLAICIYVRYF--RKKNGEEESKLPPEDSMSP---STKD---------------------------------------------------------GICTTF---------------------------------VLKFLA----------------------------------------------------------------------------------------------------------------------------------------------------------------------------------------------------------------------------------------------------------------------------------------

>Mt R108 LYK5likeKIN

----------------------------------------------------------------------------------------------------------------------------------------------------------------------------------------------------------------------------------------------MIEI----ELLKRYN--VN--FSQGSG-LVYIRGKDGNGLYVPLPP--KK-GFTF-----------------------KLIRELAILMTIYFCDLDRILVIAGISIGGSCMVLLLLLCIYVRYF--RKKNGEEESKFPPEDSMTP---STKDVDKDTNDDN--------------GSKYIWVDKSPEFSYEELANATDNFSLAKKIGQGGFGEVYYGELRGQKIAIKKMKMQATREFLSELKVLTSVHHWNLVHLIGYCVEGFLFLVYEYMENGNLSQHLHN--SE-KEPMTLSTRMKIALDVARGLEYIHDHSVPVYIHRDIKSDNILLNENFTGKVADFGLTKLTDAASSADNTD-HVAGTFGYMPPENA-YGRI--SRKIDVYAFGVVLYELISAKAAVIKIDKTEFEFKSLEIKTNE-SIDEYKSLVALFDEVMNQTGDCIDDLRKLVDPRLGYNYSIDSISKMAKLAKACINRDPKQRPTMRDVVVSLMELNSS-ID--NKNSTGSA----------------------

>Mt A17 LYK5likeKIN

----------------------------------------------------------------------------------------------------------------------------------------------------------------------------------------------------------------------------------------------MIEI----ELLKRYN--VN--FSQGSG-LVYIRGKDGNGLYVPLPP--KK-GFTF-----------------------KLIRELAILMTIYFCDLDRILVIAGISIGGSCMVLLLLLCIYVRYF--RKKNGEEESKFPPEDSMTP---STKDVDKDTNDDN--------------GSKYIWVDKSPEFSYEELANATDNFSLAKKIGQGGFGEVYYGELRGQKIAIKKMKMQATREFLSELKVLTSVHHWNLVHLIGYCVEGFLFLVYEYMENGNLSQHLHN--SE-KEPMTLSTRMKIALDVARGLEYIHDHSVPVYIHRDIKSDNILLNENFTGKVADFGLTKLTDAANSADNTV-HVAGTFGYMPPENA-YGRI--SRKIDVYAFGVVLYELISAKAAVIKIDKTEFEFKSLEIKTNE-SIDEYKSLVALFDEVMNQTGDCIDDLRKLVDPRLGYNYSIDSISKMAKLAKACINRDPKQRPTMRDVVVSLMELNSS-ID--NKNSTGSA----------------------

>Mt R108 LYK5

-------------------------MEQPLKFRLS----LLF-LL--LVLQSITSESK-CSKTCDLALASYY--IGEG-INITYISSIMQSNVVSKE----EDILSY-NTYKIKNIDAIQSDTRVNVPIPCDCI------NDEFLGHTFLYKLRL----GDIYPSIAERTYTNLTTEEWME--RVNIYPGTDLP-VSAMVNVTVNCSCGSREVSKDYGLFITYPLSSKDTLESIAKDTMIEA----ELLQRYNPGVN--FSQGSG-LVFIPGKDENGFYVPLPP--RKGH------------------------------------------LARSLGTAGISIGGLCMVLLLLLCIYVRYF--RMKNGEEKSKLSPDDSMTP---STKDVDKDTNGDT--------------GSRYIWLDKSPEFSYEELANATDNFSLAKKIGQGGFGEVYYGELRGQKIAIKKMKMQATREFLSELKVLTSVHHRNLVHLIGYCVEGFLFLVYEYMENGNLNQHLHN--SE-KEAITLSTRMKIALDVARGLEYIHDHSIPVYIHRDIKSDNILLNENFTGKVADFGLTKLTDAASSADNTD-HVAGTFGYMPPENA-YGRI--SRKIDVYAFGVVLYELISAKAAVIKIDKTEFELKSLEIKTNE-SIDEYKSLVALFDEVMDQTGDPIEGLRKLVDPRLGYNYSIDSISKMAKLAKACINRDPKQRPKMRDVVVSLMKLNYT-ID--DESRTGSAELSLAVEHDSN-----------

>Mt A17 LYK5

-------------------------MEQPLKFRLS----LLF-LL--LVLQSITSESK-CSKTCDLALASYY--IRPG-TTLANISKVMQSNVVSKE----EDILSY-NTA-ITNIDAIQSDTRVNVPFPCDCI------NDEFLGHTFLYKLRL----GDIYPSIAERTYTNLTTEEWME--RVNSYPGTDLP-VSAMVNVTVNCSCGSREVSKDYGLFITYPLSSKDTLESISKDTMIEA----ELLQRYNPGVN--FSQGSG-LVFIPGKDENGFYVPLPP-----S------------------------------------------L----GTAGISIGGLCMVLLLLLCIYVRYF--RMKNGEEKSKLSPDDSMTP---STKDVDKDTNGDT--------------GSRYIWLDKSPEFSYEELANATDNFSLAKKIGQGGFGEVYYGELRGQKIAIKKMKMQATREFLSELKVLTSVHHRNLVHLIGYCVEGFLFLVYEYMENGNLNQHLHN--SE-KEPITLSTRMKIALDVARGLEYIHDHSIPVYIHRDIKSDNILLNENFTGKVADFGLTKLTDAASSADNTD-HVAGTFGYMPPENA-YGRI--SRKIDVYAFGVVLYELISAKAAVIKIDKTEFELKSLEIKTNE-SIDEYKSLVALFDEVMDQTGDPIEGLRKLVDPRLGYNYSIDSISKMAKLAKACINRDPKQRPKMRDLVVSLMKLNYT-ID--DESRTGSAELSLAVEHDSN-----------

>Mt A17 LYK8

---------------------------MITNQIFI----LNFLLFVLLIMKTVRS----CNSGCDLALASYY--IEEG-TNLTYISNLFNQPT--------SEILKY-NPN-IQNPDTIQSHTRLNIPFTCDCL------SGLFLGHTFSYKLKE----GENYKAVANGYYSNLTTIDFLI--RVNSYPATDIP-AGTVINVTVNCSCGDRDVSKDYGLFLTYPLRNGDSLPGIAMESGVPV----ELVKRYNPASN--FRAGE--LVFLPAKDENGNFPPLKM--GS-G------------------------------------------MSKG-GIVGIVVGGAFGILLLVLILYVVFY--RRKKVADQVTLLPVPGASE---LDQSSQLQHGRGSSMDKTSESTTVVSPRLTGITVDKSVEFSYEELAKATDGFSTANIIGRGGFGLVYYAELRNEKAAIKKMDMQASKEFLAELKVLTHVHHLNLVRLIGYCVEGSLFLVYEYIENGNLSQHLRG--TG-KDPLSWPARVQIALDSARGLEYIHEHTVPVYIHRDVKSANILIDKNFRGKVADFGLTKLTEYGSSSLQT--RLVGTFGYMPPEYAQYGEV--SPKIDVYAFGVVLFELISGKQAI--------------VKTDE-AKNESKGLVALFEEVLGLS-EPKEDLGKLVDPRLGENYPIDSVFKMSQLAKACTHENPQLRPSMRSIVVALMTLSSA-AE--DWDVGSFYENQALVHLMSGR----------

>Mt R108 LYK8

---------------------------MITNQIFI----LNFLLFVLLIMKTVRS----CSSGCDLALASYY--IEEG-TNLTYISNLFNQPT--------SEILKY-NPN-IKSPDTIQSNTRLNIPFTCDCL------SGLFLGHTFSYKLKE----GENYKAVANGYYSNLTTIDFLI--RVNSYPATDIP-AGTVINVTVNCSCGDRDVSKDYGLFLTYPLRNGDSLPGIAMESGVPV----ELVKRYNPASN--FRAGE--LVFLPAKDENGNFPPLKM--GS-G------------------------------------------MSKG-GIVGIVVGGAFGILLLVLILYVVFY--RRKKVADQVTLLPVPGASE---LDQSSQLQHGRGSGMDKTSESTTVVSPRLTGITVDKSVEFSYEELAKATDGFSTANIIGRGGFGLVYYAELRNEKAAIKKMDMQASKEFLAELKVLTHVHHLNLVRLIGYCVEGSLFLVYEYIENGNLSQHLRG--TG-KDPLSWPARVQIALDSARGLEYIHEHTVPVYIHRDVKSANILIDKNFRGKVADFGLTKLTEYGSSSLQT--RLVGTFGYMPPEYAQYGEV--SPKIDVYAFGVVLFELISGKQAI--------------VKTDE-AKNESKGLVALFEEVLGLS-EPKEDLGKLVDPRLGENYPIDSVFKMSQLAKACTHENPQLRPSMRSIVVALMTLSSA-AE--DWDVGSFYENQALVHLMSGR----------

>Glycine max LYK1

-----------------------MTTHPTTKSKPP---HVFFLLLIQLLISITRVKGS-CVTGCNLALASYY--LGNG-TNLTYISNLFGRPT--------SEILKY-NPS-VKNPNVILSQTRINVPFSCDCL------NGAFLGHTFSYAIQH----GNTYKIVAEVDFSNLTTEDWVG--RVNSYPPNQIP-DNVNINVTVNCSCGNRHVSKDYGLFMTYPLRVGDSLQRVAAEAGVPA----ELLLRYNPTAD--FGAGNG-LVFVPAKDENGNFPPMQL--RS-G------------------------------------------ISSG-AIAGIAVGGAVGVLILALLLYVGLR--RRRKV-AEVSLLPVPGASEDQCSPLQLHHGIGCGSSLDKASESSVVASPRLTGITVDKSVEFPYEELDKATDGFSAANIIGRGGFGSVYYAELRNEKAAIKKMDMQASNEFLAELNVLTHVHHLNLVRLIGYCVEGSLFLVYEYIENGNLSQHLRG--SG-RDPLTWAARVQIALDAARGLEYIHEHTVPVYIHRDIKSANILIDKNFRAKVADFGLTKLTEYGSSSLHT--RLVGTFGYMPPEYAQYGDV--SSKIDVYAFGVVLYELISGKEAI--------------VRTNE-PENESKGLVALFEEVLGLS-DPKVDLRQLIDPTLGDNYPLDSVFKVSQLAKACTHENPQLRPSMRSIVVALMTLSSA-TE--DWDVGSFYENQALVHLMSGR----------

>Lotus japonicus LYS7

-----------------------MTTKPNRAFFLI----YFLFLL----LIIIKAQGS-CVSGCNLALASYT--IWQG-ANLTYISKLFGKEP--------SEIMKY-NPN-VKNPDVIQSETQINVPFSCECL------DGIFQGHTFSYTMQA----GNTYKSIAKVDFSNLTTEEWVT--RVNSYKPNDIP-IGVKINVTINCSCGDERVSKGYGLFLTYPLRPGDDLPRLAVESGVSA----EVLQGYNAGAD--FSAGNG-LVFLPAKDENGNFPPLQKLGRS-G------------------------------------------ISPG-AIAGIVVGGAVVILLLAFASYVGLN--RRTKV-DEVSLLPVPGSYE---DHNSQQLHHGCGSSMYKASESSTVVSPRLTGITVDKSVEFPYEELAKATDSFSNANIIGRGGFGSVYYAELRNEKAAIKKMDMQASNEFLAELKVLTHVHHLNLVRLIGYCVEGSLFLVYEYIENGNLSEHLRG--SG-RDPLSWPARVQIALDSARGLEYIHEHTVPVYIHRDIKSANILIDKNFRGKVADFGLTKLTEYGSSSLQT--RLVGTFGYMPPEYAQYGEI--SPKVDVYAFGVVLYELVSGKEAI--------------VRTNG-PENESKALIALFEEVLGQP-DPKEYLGKLVDPRLGDSYPLDSVFKVSQLAKACTHENPQLRPSMRSIVVALMTLTCA-AE--DWDVGSFYENQALVHLMSGR----------

>Populus trichocarpa LYK12

-------------------MIPSSSRYPHIQTLLV----SCVLLF-----LVFKVQAK-CRTGCGLALASYY--VWQG-SNLTYISTIFNQSI--------TEILRY-NPK-VPNQDSIRSDTRLNVPFSCDCL------NGDFLGHTFSYITQS----GDTYHKIARNAFSNLTTEDWVH--RVNIYDITEIP-NYVPINVTVNCTCGDKQVSRDYGLFTTYPLRPDENLSSLEAESGVPA----DLLEKYNLGTD--FNAGGG-IVYMPAKDPTGNYPPLKI--AT-G------------------------------------------ISSR-AIAGISVAGVAGSFFLASCFYFGFY--RRREV--EASLFPEAAESP---YIHHRHGSGNILEQTSETAALVG--SPGLTGFTVDKSVEFSYEELAKATNDFSMDNKIGQGGFGAVYYAELRGEKAAIKKMDMQASKEFLAELKVLTHVHHLNLVRLIGYCVEGSLFLVYEFIENGNLGQHLRS-NSG-KDPLPWSTRVQVALDSARGLEYIHEHTVPVYIHRDVKSANILIDKNFRGKVADFGLTRLTEVGSASLHT--RLVGTFGYMPPEYAQYGDV--SSKIDVYAFGVVLYELISAKEAV--------------VKTNE-FITESMGLVALFEEVLGQP-DPRENLPKLVDARLGDDYPLDSVCKMAQLARACTQENPHVRPSMRSIVVALMTLSSS-TE--DWDVGSLYENQAIVDLMSGR----------

>Solanum lycopersicum LYK12

----------------------MNIRAKTINLSSF----FLFIIL--------NGEAKSCGNGCEMAIASYH--IWSG-ANLTYISHLFNLTI--------PVILNY-NPQ-ITNQDSITSDTRINLPFSCDCL------NGDFLGHTFVYKTVF----GDTYKKVATMAFANLTTEYWLK--RVNNYDPTSIP-DYAMINVTVNCSCGDGEVSDDYGLFATYPIRPGENLSTVAVGSGVPA----ELLQKFNPGLD--FGSGSG-IVFVPARDAHGNFPPLKT--RSRG------------------------------------------LSRG-AIAGTTVAAIFGATFFVVCVYFVFY--RSKQA-EEESFLQGSSD-----EHFNENFRPPNLEKITESGPLFGVISPRPTGITVDKSVEFSYEELAKATNNFSMENKIGQGGFGLVFYGMLKGERAAIKKMDMQASKEFFAELKVLTHVHHLNLVRLIGYCVEGSLFLVYEYIENGNLGEHLRG--SS-RNPLSWSTRVQIALDAARGLEYIHEHTVPLYIHRDIKSANILIDKDFRAKVADFGLTKLTEVGSTSFHT--RLVGTFGYMPPEYAQYGDV--SPKVDVYAFGVVLYELISAKEAI--------------VKTNE-VITESKGLVALFEDVLHQSGGAREGLCKVVDPKLGDDYPLDSVCKVAQLAKACTHENPQLRPSMRSIVVALMTLSSS-TE--DWDIGSFYEN----HLMSGR----------

>Mt A17 LYK6

---------------------------MKISFSFI----VLILLI-------ASTESK-CNEGCSLALASYT--LNHV-SNLTYISNIMKSNVLSKP----QDIIINNDKN-----------KRANVPFPCNCI------NGEFLAYTFLYELQP----GETYTSVAEESFSNLTTDVWMQ--NFNVYRPTNIP-DFAMIKVTVNCSCGNKEVSMDYGLFITYPLRSEDTLESIAKGAEIEA----ELLQRYNPGVN--FSKGSG-LVFIPGKDQNGSYLPLHP--STVG------------------------------------------LGTV-AITGISVGVLAALLLLLFFVYIKYYLKKKNKKTWEKNLILDDSKMK---SAQIGT---------------------NIASIMVEKSEEFSYKELSIATNNFSMANKIGEGGFGEVFYAELRGQKAAIKKMKMKASKEFCAELKVLTLVHHLNLVGLIGYCVEGFLFLVYEYIDNGNLSQNLHD--SE-REPLSWSTRMQIALDSARGLEYIHEHTVPVYIHRDIKSENILLDKSFCAKVADFGLSKLADVGNSTSSTI-VAEGTFGYMPPEYA-CGSVSSSPKVDVYAFGVVLYELISAKAAV--------------INDGP----QVTGLVAVFDEVFGYDQDPTEGIKNLVDPRLGDNYSIDSVCKMAQLAKACTMRDPQLRPSMRSIVVALMTLTST-TE--DWNISSFYENPAFLNLMSGK----------

>Mt R108 LYK6

---------------------------MKISFSFI----VLILLI-------ASTESK-CNEGCSLALASYT--LNHV-SNLTYISNIMKSNVLSKP----QDIIINNNKN-----------KRVNVPFPCDCI------NGEFLAYTFLYELQP----GETYTSVAEESFSNLTTDVWMQ--NFNVYRPTNIP-DFAMIKVTVNCSCGNREVSMDYGLFITYPLSSKDTLESIAKDTKIEA----ELLQRYNPGVN--FSKGSG-LVFIPGKDQNGSYLPLHP--STVG------------------------------------------LGTV-AITGISVGVLAALLLLLFFVYIKYYLKKKNKKTWEKNLILDDSKMK---SAQIGT---------------------NIASIMVEKSEEFSYKELSIATNNFSIANKIGEGGFGEVFYAELRGQKAAIKKMKMKASKEFCAELKVLTLVHHLNLVGLIGYCVEGFLFLVYEYIDNGSLSQNLHN--SE-REPLSWSTRMQIALDSASGLEYIHEHTVPVYIHRDIKSENILLDKSFCAKVADFGLSKLADVGNSTSSTI-VAEGTFGYMPPEYA-CGSVSSSPKVDVYAFGVVLYELISAKAAV--------------INDGP----QVTGLVAVFDEVFGYDQDPTEGIKNLVDPRLGDNYSIDSVCKMAQLAKACTMRDPQLRPSMRSIVVALMTLTST-TE--DWNISSFYENPAFLNLMSGR----------

>Lotus japonicus NFRe

-------------------------MEPKLTFSLS----FLLTLL------SPFAESK-CIKGCDLALASYY--QWSG-SNLTYISKIMESQILSKP----QDIVTY-NKG---KRNFGVFSTRVNVPFPCDCI------NGEFLGHTFEYQLQP----EETYTTVASETFSNLTVDVWMQ--GFNIYPPTNIP-DFAVLNVTVNCSCGNSEVSKDYGLFITYPLRIEDSLQSIAEEMKLEA----ELLQRYNPGVN--FSQGSG-LVFIPGKDQNGSYVPFQQ--STVG------------------------------------------FSGG-VIAGISVGVLVGLLLVAFCVYTKHL--QKKKA-LEKKLILDDSTVN---SAQVSNDSG---------------------GIMMDKSREFSYKELADATNNFSVANRIGEGGFGTVYYADLSGEKTAIKKMNMLASREFLAEVKVLANVHHLNLVRLIGYCIEGSLFLVYEYIDNGNLKQSLHD--LE-REPLPWSTRVQIALDSARALEYIHEHTVHVYIHRDIKSENILLDNSFHAKVADFGLSKLVQVGNSIGSSVNMMKGTFGYMPPEYA-RGVVSPSPKIDVYAFGVVLYELISAKEAV--------------IRDGA----QSKGLVALFDEVLGNQLDPRESLVSLVDPRLQDNYSIDSVCKMAQLAKVCTERDPTGRPSMRSVMVALMTLSST-TQ--SWDIASFYENPALVNRMSGRLE--------

>Solanum lycopersicum LYK1

----------------------MFESRPRSVLSLG----VFV-ILVYLSSVPLPVNSQ-CNRGCDLALASFY--VWRG-SNLTLISEMFSTSI--------ADIVSYNNRDNIPNQDSVIAGTRINIPFRCDCLN-----DGEVLGHAFPYRVKS----GDTYDLVAR-NYSDLTTAQWMM--KFNSYPENNIP-NTVNLSVVVNCSCGNSDVSKDFGLFVTYPVRAEDNLTSVASAANVSE----DIIRRYNPAAVSILDIGQG-IIYIPGRDRNGNFPPLPT--STDG------------------------------------------LSGG-AKAGISIGAIGVVLLLAGLVYVGCY--RNKTR-KISLLRSEDHL-----HQYGHGPEGSTTVKAADSGRLADGNSPVLSGITVDKSVEFTYEELATATNDFSIANKIGQGGFGAVYYAELRGEKAAIKKMDMEATREFLAELKVLTNVHHLNLVRLIGYCVEGSLFLVYEYVENGHIGQHLRG--TG-RDPLPWSKRVQIALDSARGLEYIHEHTVPVYIHRDIKTANILIDKNFHAKVADFGLTKLTEVGSSSLQT--RLVGTFGYMPPEYAQYGDV--SPKVDVYAFGVVLYELISAKEAI--------------VKPNG-SVTESKGLVALFEEVLNQP-DPDEDLRQLVDPRLGDDYPLDSVRKMAQLAKACTHENPLIRPSMRSIVVALMTLSSS-TE--DWDVGSFYGNQGMINLMSGR----------

>Glycine max LYK4

---------------------------MEHSFRLP----VFF-LL--CASIAFSAESK-CSRGCDLALASYY--LSQG--DLTYVSKLMESEVVSKP----EDILSY-NTDTITNKDLLPASIRVNVPFPCDCI------DEEFLGHTFQYNLTT----GDTYLSIATQNYSNLTTAEWLR--SFNRYLPANIP-DSGTLNVTINCSCGNSEVSKDYGLFITYPLRPEDSLQSIANETGVDR----DLLVKYNPGVN--FSQGSG-LVYIPGKDQNAIYVPLHL--SSGG------------------------------------------LAGG-VIAGISIGVVTGLLLLAFCVYVTYY--RRKKV-WKKDLLSEESRKN---SARVKNDEASGDSAAEGGT--------NTIGIRVNKSAEFSYEELANATNNFSLANKIGQGGFGVVYYAELNGEKAAIKKMDIQATREFLAELKVLTHVHHLNLVRLIGYCVEGSLFLVYEYIENGNLGQHLRK--SG-FNPLPWSTRVQIALDSARGLQYIHEHTVPVYIHRDIKSENILIDKNFGAKVADFGLTKLIDVGSSSLPTV-NMKGTFGYMPPEYA-YGNV--SPKIDVYAFGVVLYELISGKEAL--------------SRGGV-SGAELKGLVSLFDEVFDQQ-DTTEGLKKLVDPRLGDNYPIDSVCKMAQLARACTESDPQQRPNMSSVVVTLTALTST-TE--DWDIASIIENPTLANLMSGK----------

>Mt A17 LYK7

--------------------------MKPIKFRLS----FLF-ML--LASKSFIAESK-CSKTCNIALASYY--LQDD-TNLTYVSNIMQSNLVTKP----EDIVSY-NTDTITNKDFVQSFTRVNVPFPCDCI------HDEFLGHIFQYQVAT----KDTYLSVASNNYSNLTTSEWLQ--NFNSYPSNDIP-DTGTLNVTVNCSCGNSDVSKDYGLFITYPLRPEDSLELISNKTEIDA----ELLQKYNPGVN--FSQGSG-LVYIPGKDQNRNYVPFHI--STGG------------------------------------------LSGG-VITGISVGAVAGLILLSFCIYVTYY--RKKKI-RKQEFLSEES------SAIFGQVKNDEVSGNATYGTSDSASPANMIGIRVEKSGEFSYEELANATNNFNMANKIGQGGFGEVYYAELNGEKAAIKKMDMKATKEFLAELKVLTRVHHVNLVRLIGYCVEGSLFLVYEYIDNGNLGQHLRS--SD-GEPLSWSIRVKIALDSARGLEYIHEHTVPTYIHRDIKSENILLDKNFCAKVADFGLTKLIDAGISSVPTV-NMAGTFGYMPPEYA-YGSV--SSKIDVYAFGVVLYELISAKAAV--------------IMGED-SGADLKGLVVLFEEVFDQP-HPIEGLKKLVDPRLGDNYPIDHVFKMAQLAKVCTNSDPQQRPNMSSVVVALTTLTST-TE--DWDITSIFKNPNLVNLMSGR----------

>Mt R108 LYK7

--------MIHHTPKFLSYVPIILYTMKPIKFRLS----FLF-ML--LASKSFIAESK-CSKTCDIALASYY--LQDD-TNLTYVSNIMQSNLVTKP----EDIVSY-NTDTITNKDFVQSFTRVNVPFPCDCI------HDEFLGHIFQYQVAT----KDTYLSVASNNYSNLTTSEWLQ--NFNSYPSNDIP-DTGTLNVTVNCSCGNSDVSKDYGLFITYPLRPEDSLELISNKTEIDA----ELLQKYNPGVN--FSQGSG-LVYIPGKDQNRNYVPFHT--STGG------------------------------------------LSGG-VITGISVGAVAGLMLLSFCIYVTYY--RKKKI-RKQEFLSEESSAIFG-QVKNDEVSGNATYGTSDSASPA-----NMIGIRVEKSGEFSYEELANATNNFNMANKIGQGGFGEVYYAELNGEKAAIKKMDMKATKEFLAELKVLTRVHHVNLVRLIGYCVEGSLFLVYEYIDNGNLGQHLRS--SD-GEPLSWSTRVKIALDSARGLEYIHEHTVPTYIHRDIKSENILLDKNFCAKVADFGLTKLIDAGISSVPTA-NMAGTFGYMPPEYA-YGSV--SSKIDVYAFGVVLYELISAKAAV--------------IMGGD-SGADLKGLVVLFEEVFDQP-HPIEGLKKQVDPRLGDNYPIDHVFKMAQLAKVCTNSDPQQRPNMSSVVVALTTLTST-TE--DWDITSIFKNLNLVNLMSGR----------

>Populus trichocarpa LYK1

-------------------------MNPKLGFGFL----LLL-LL------CYSIDSK-CSKGCDLALASYY--VWQG-ANLSFIAEVMQSSILKSTDF--DTILRY-NPQ-VTNKDSLPSFIRISIPFPCECI------NGEFLGHFFTYNVRS----QDTYGTVADTYYANLTTTPSLI--NFNSYPEVNIP-DNGVLNVSVNCSCGDSSVSKDYGLFMTYPLRPNDTLASIANQTNLTQ----SLLQRYNVGFD--FNQGSG-VVYIPTKDPDGSYLPLKS--ST-G------------------------------------------IAGG-VVAGICIAAVAVALLLAVFIYVGFY--RKKKV-KGAILLPASQEL----SPRIVQVPGSNSNKPVDATGFQ-----GLTGLTVDKSVVFSYEELAKATDDFSLANKIGQGGFGSVYYAELRGEKAAIKKMDMQASKEFLAELKVLTHVHHLNLVRLIGYCVEGSLFLVYEFIENGNLSQHLRG--SE-KDPLPWSTRVQIALDSARGLEYIHEHTVPVYIHRDIKSANILIDKNFRGKVADFGLTKLTEVGSTSLPT--RLVGTFGYMPPEYAQYGDV--SPKVDVYALGVVLYELISAKEAI--------------VKSNG-SSAESRGLVALFEDVLNQP-DPREDLRKVVDPRLGEDYPLDSVRKMAQLGKACTQENPQLRPSMRSIVVALMTLSSS-TE--DWDVGSFYENQALVNLMSGR----------

>Populus trichocarpa LYK11

-------------------------MNPKLGLGFI----LLL-LL------CYSIESK-CRKGCDLALASYY--VWQD-ANLTFIAEVMQSSILKSSDF--DTILRY-NPQ-LPSKDSLSSLIRINIPFPCDCI------EGQFLGHFFNFNVRS----QNTYTVVADTYYAKLTTIPSLM--YFNNYSEFNIP-DNGKLNVSVNCSCGDSSVSKDYGLFMTYPLQPNDTLNSIANQTNVTQ----ELLQRYNVGFN--FSRGTG-VVYIPTKDADGSYRPLKS--ST-G------------------------------------------IAGG-AIAGISIAAVAVALLLAVLIYVGFY--RKKK--EKGAILLSASPQL---SPRILHVTGSNTPVNATGSQ-------GLAGITVDKSVEFSYEELAKATDDFSFANKIGEGGFGTVYYAELRGEKAAIKKMDVQDSKEFFAELKVLTHVHHLNLVRLIGYCVEGSLFVVYEYIENGNLSQHLRG--SG-KDPLTWSTRVQIALDSARGLEYIHEHTVPVYIHRDIKSANILIDKNFRGKVADFGLAKLTKVGSASLLT--RLVGTFGYMSPEYAQYGDV--SPKLDVFAFGVVLYELISAKEAI--------------VKAND-SSAESRGLIALFENVLNQP-DPGEDLRKLVDPRLGEDYPLDSVRKVTQLAKACTHENPQMRPSMRSIVVALMTLSSS-TE--DWDVGSFYENKALVNLMSGR----------

>Lupinus angustifolius LYK3

-------------------------MEPIFWFLIK----LSPLFF-----LCSNAESK-CTQGCPIALASYY--MLSG-SNLTYISQIMSSHVLHSP----EDIVSY-------NKDKVQPFTRVNVPFPCDCI------KGEFLGHMFQYVVQT----GDTYETVAGTNYANLTNVEWLR--RFNTYLPDNIS-STGMLNVTVNCSCGNSDVS-DYELFITYPLRPGETLGSVAKSVKLDS----GLLQRYNPSVN--FNQGSG-LVYIPGKDQNGSYVFLSS--SSGG------------------------------------------LAGG-AIAGIAVGVVAGILLLVVCIYVGCF--RKKKI-QKEEVVRPDSKSH---SVPDGMDEISLVAAYETSRPRGSA---AIAGISMDKSVEFSYEELASATNNFSVANKIGQGGFAVVYYAELRGEKAAIKKMDMQASKEFLAELNVLTHVHHLNLVRLIGYSIKGSLCLVYEFIENGNLSQHLHG--SG-REPLPWTIRVQIALDSARGLEYIHEHTMPIYIHRDIKSANILIDKNFRGKVADFGLAKLAEVGSSLRPTV-RLVGTFGYMPPEYAQYGDV--SPKVDVYAFGVVLYELISAKEAI--------------IQ----SIVDSKGLVHWFKEVLSQP-HSTEDLCKLVDPKLGDNYPIDSVLKLAQLAKACTQHNPQLRPSMRSIVVALMTLSST-TD--DWDVGSFYENQNLVNLMSGK----------

>Lupinus angustifolius LYK4

-------------------------MKPILMFLIM----FLLWLL-----LLSSAESK-CTQGCSLALASYY--MYSG-STLTSISQVMSSQLLQIP----EDIVTY-NKDTIPNKDSVQAFIRVNVPFPCDCI------DGEFLGHMFQYDVKT----GDTYQLVAETEYANLTNIDWLM--KFNSYPANNIP-DTGTLNVTVNCSCGEKNVS-NYGLFITYPLRPGDTLDSVSKSVDLDS----GLLQRYNPGVN--FNQGSG-LVYIPGKDQNGSYVFLNS--SSEG------------------------------------------LAGG-VIAGIVIGVLAGILLLVAGIYVGYF--RKKKI-QKEELLEQDSKSL---FVQNGMDETARTA--------------ATTGISVDKSVEFSYEELASATDNFSMANKIGQGGFGVVYYAELRGEKAAIKKMDMQASKEFLAELKVLTHVHHLNLVRLIGYSIEGSLFLVYEFIENGNLSQHLRG--SG-RDPLPWPARVQIALDSARGLEYIHEHTVPVYIHRDIKSANILIDKNFRGKVADFGLTKLTEVGSSSLPTG-RLVGTFGYMPPEYAQYGDV--SPKVDVYAFGVVLYELISAKEAI--------------IQANE-SIADSKGLVALFEGVLNQP-DSTEDLCKVVDPRLGDNYPIDSVRKLAQLAKACTQDNPQLRPSMRSIVVALMTLSST-TD--DWDVGSFYENQNLVNLMSGR----------

>Glycine max LYK2

--------------------------MEALRLAYL----LLPWWL-----VFSTAESA-CKEGCGVALGSYY--LWRG-SNLTYISSIMASSLLTTP----DDIVNY-NKDTVPSKDIIIADQRVNVPFPCDCI------DGQFLGHTFRYDVQS----QDTYETVARSWFANLTDVAWLR--RFNTYPPDNIP-DTGTLNVTVNCSCGNTDVA-NYGLFVTYPLRIGDTLGSVAANLSLDS----ALLQRYNPDVN--FNQGTG-LVYVPGKDQNGSFVRLPS--SSGG------------------------------------------LTGR-AIAGIAVGIVAALLLLGVCIYVGYF--RKKI--QKDEFLPRDSTAL---FAQDGKDETSRSSANETSGPGGPA---IITDITVNKSVEFSYEELATATDNFSLANKIGQGGFGSVYYAELRGEKAAIKKMDMQASKEFLAELNVLTRVHHLNLVRLIGYSIEGSLFLVYEYIENGNLSQHLRG--SGSREPLPWATRVQIALDSARGLEYIHEHTVPVYIHRDIKSANILIDKNFRGKVADFGLTKLTEVGSSSLPTG-RLVGTFGYMPPEYAQYGDV--SPKVDVYAFGVVLYELISAKEAI--------------VKTND-SVADSKGLVALFDGVLSQP-DPTEELCKLVDPRLGDNYPIDSVRKMAQLAKACTQDNPQLRPSMRSIVVALMTLSST-TD--DWDVGSFYENQNLVNLMSGR----------

>Mt A17 LYK9

-----------------------MEHQPRFTSFIS---LPLFSIF--LASIPFITESK-CTKGCSLALANFY--VSQG-SNLTYISSIMRSNIQTRP----EDIVEY-SREIIPSKDSVQAGQRLNVPFPCDCI------DGQFLGHKFSYDVET----GDTYETVATNNYANLTNVEWLR--RFNTYPPNDIP-DTGTLNVTVNCSCGDADVG-NYALFVTYPLRPGETLVSVANSSKVDS----SLLQRYNPGVN--FNQGSG-IVFVPGKDQNGSFVFLGS--SS-G------------------------------------------LGGG-AIGGIAVGIVVVLLLVAAAIYFGYF--RKKKI-QKEELFSRDSTAL---FSQDGKDENSHGAANVTQRPG------VMTGITVDKSVEFSYDELAAASDNFSMANKIGQGGFGSVYYAELRGEKAAIKKMDMQATKEFLAELKVLTRVHHLNLVRLIGYSIEGSLFLVYEYIENGNLSQHLRG--SG-RDPLPWATRVQIALDSARGLEYIHEHTVPVYIHRDIKPANILIDKNFRGKVADFGLTKLTEVGSSSLPTG-RLVGTFGYMPPEYAQYGDV--SPKVDVYAFGVVLYELISAKEAI--------------VKSSE-SVADSKGLVGLFEGVLSQP-DPTEDLRKIVDPRLGDNYPADSVRKMAQLAKACTQENPQLRPSMRSIVVALMTLSST-TD--DWDVGSFYENQNLVNLMSGR----------

>Mt R108 LYK9

-----------------------MEHQPRFTSFIS---LPLFSIL--LASIPFITESK-CTKGCSLALANFY--VSQG-SNLTYISSIMRSNIQTRP----EDIVEY-SREIIPSKDSVQAGQRLNVPFPCDCI------DGEFLGHKFSYDVET----GDTYETVATNNYANLTNVEWLR--RFNTYPPNDIP-DTGTLNVTVNCSCGDSDVG-NYALFVTYPLRPGETLGSVANSSKVDS----SLLQRYNPGVN--FNQGSG-IVFVPGKDQNSSFVFLGS--RS-G------------------------------------------LGGG-AIGGIAVGIVVVLLLVAAAIYFGYF--RKKKI-RKEELFSRDSTAL---FSQDGKDENSHGAANVTQRPG------VMTGITVDKSVEFSYDELAAASDNFSMANKIGQGGFGSVYYAELRGEKAAIKKMDMQATKEFLAELKVLTRVHHLNLVRLIGYSIEGSLFLVYEYIENGNLSQHLRG--SG-RDPLPWATRVQIALDSARGLEYIHEHTVPVYIHRDIKPANILIDKNFRGKVADFGLTKLTEVGSSSLPTG-RLVGTFGYMPPEYAQYGDV--SPKVDVYAFGVVLYELISAKEAI--------------VKSSE-SVADSKGLVGLFEGVLSQP-DPTEDLRKIVDPRLGDNYPADSVRKMAQLAKACTQENPQLRPSMRSIVVALMTLSST-TD--DWDVGSFYENQNLVNLMSGR----------

>Lotus japonicus CERK6

------------------------MEHPRLGFPIT----LLLFSF---ILLPSTSQSK-CTHGCALAQASYY--LLNG-SNLTYISEIMQSSLLTKP----EDIVSY-NQDTIASKDSVQAGQRINVPFPCDCI------EGEFLGHTFQYDVQK----GDRYDTIAGTNYANLTTVEWLR--RFNSYPPDNIP-DTGTLNVTVNCSCGDSGVG-DYGLFVTYPLRPGETLGSVASNVKLDS----ALLQKYNPNVN--FNQGSG-IVYIPAKDQNGSYVLLGS--SSGGLTFFSEIWLWKDCSHIHVEHMVSLYKKGDADLSSYFASIYAGLAGG-AIAGIAAGVAVCLLLLAGFIYVGYF--RKKRI-QKEELLSQETRAI---FPQDGKDENPRSTVNETPGPGGPA---AMAGITVDKSVEFSYDELATATDNFSLANKIGQGGFGSVYYAELRGERAAIKKMDMQASKEFLAELKVLTRVHHLNLVRLIGYSIEGSLFLVYEFIENGNLSQHLRG--SG-RDPLPWATRVQIALDSARGLEYIHEHTVPVYIHRDIKSANILIDKNYRGKVADFGLTKLTEVGSSSLPTG-RLVGTFGYMPPEYAQYGDV--SPKVDVYAFGVVLYELISAKDAI--------------VKTSE-SITDSKGLVALFEGVLSQP-DPTEDLRKLVDQRLGDNYPVDSVRKMAQLAKACTQDNPQLRPSMRSIVVALMTLSST-TD--DWDVGSFYENQNLVNLMSGR----------
